# Comparative genomics of three *Colletotrichum scovillei* strains and genetic analysis revealed genes involved in fungal growth and virulence on chili pepper

**DOI:** 10.1101/2021.09.06.459084

**Authors:** Dai-Keng Hsieh, Shu-Cheng Chuang, Chun-Yi Chen, Ya-Ting Chao, Mei-Yeh Jade Lu, Ming-Che Shih, Miin-Huey Lee

**Affiliations:** Ph.D. Program in Microbial Genomics, National Chung-Hsing University and Academia Sinica, Taichung, 40227, Taiwan; Agricultural Biotechnology Research Center, Academia Sinica, Taipei 11529, Taiwan; Biodiversity Research Center, Academia Sinica, Taipei 11529, Taiwan; Departmment of Plant Pathology, National Chung-Hsing University, Taichung 40277, Taiwan; Advanced Plant Biotechnology Center, National Chung-Hsing University, Taichung 40277, Taiwan

## Abstract

*Colletotrichum scovillei* is a virulent pathogen and the dominant species causing anthracnose of chili pepper in many Asian countries. Three strains of this pathogen, Coll-524, Coll-153 and Coll-365, show varied virulence on chili pepper fruit. Among the three strains, Coll-365 showed significant defects in growth and virulence. To decipher the genetic variations among these strains and identify genes contributing to growth and virulence, in this study, comparative genomic analysis and gene transformation to verify gene function were applied. The genomes of the three strains were sequenced and Coll-524 had 1.3% and 1.5% more genes than Coll-153 and Coll-365, respectively. Compared to Coll-524 and Coll-153, Coll-365 had numerous gene losses including 33 effector genes that are distributed in different scaffolds and a cluster of 14 genes in a 34-kb genomic fragment. Through gene transformation, three genes in the 34-kb fragment were identified to have functions in growth and/or virulence of *C. scovillei.* Gene 15019 encoding a protein related to phospholipase A2-activating protein enhanced the growth of Coll-365. A combination of 15019 with one transcription factor gene 15022 and one C6 zinc finger domain-containing protein gene 15029 was found to enhance the pathogenicity of Coll-365. Introduction of gene 15215, which encodes a LysM domain-containing protein, into Coll-365 caused a reduction in the germination rate of Coll-365. In conclusion, the higher virulent strain Coll-524 had more genes and encoded more pathogenicity related proteins and transposable elements than the other two strains, which may contribute to the high virulence of Coll-524. In addition, the absence of the 34-kb fragment plays a critical role in the defects of growth and virulence of strain Coll-365.

**Author Summary:** *Colletotrichum scovillei* is a highly virulent and dominant pathogen causing anthracnose of chili that leads to significant economic loss in chili production in many Asia countries. In this study we focus on finding the gene differences of three *C. scovillei* strains with different pathogenicity in chili pepper infection and verifying the function of some genes in the lowest virulence strain. We sequenced them and did gene annotation and genome comparison. We setup a simple mathematical method to identify gene variations between strong and weak virulence strains. Our results show that the lowest virulence strain has less pathogenicity-related genes. We also found that the absence of 14 genes in a compact genomic fragment was part of the reason of growth and virulence defect of the lowest virulence strain. We identified four genes that play roles on fungal growth and/or virulence on chili pepper. We also found a group of effector genes that specifically appear in species collected form infected chili in *C. acutatum* species complex. Our research provides detailed information for why the three strains have different virulence on chili pepper.

## Introduction

Chili pepper (*Capsicum* spp.) is a globally significant spice crop. The cultivation of chili pepper is frequently threatened by the attacks of various pathogens, especially the anthracnose pathogens of the *Colletotrichum* species. *Colletotrichum* contains a large number of species members that have been classified into several species complexes [1]. *Colletotrichum* pathogens associated with *Capsicum* plants come from diverse species complexes including *C. acutatum*, *C. boninense*, *C. gloeosporioides*, *C. orchidearum* and *C. truncatum* complexes [2–6]. At least 28 species have been reported and most of them are from the acutatum, gloeosporioides and truncatum complexes [7, 8].

*C. scovillei* (formerly *C. acutatum*) is a highly virulent and dominant pathogen causing anthracnose of chili that leads to significant economic loss in chili production in many Asia countries [4, 8]. It can infect many different species of *Capsicum*, especially species that are mainly grown for human consumption such as *C. annuum*, *C. frutescens*, *C. chinense*, and *C. baccatum* [4, 8, 9]. *C. scovillei* is the most dominant species causing anthracnose of chili in Asia, and is found widely distributed in Indonesia, Malaysia, Thailand and Taiwan [8, 10]. Two pathotypes of *C. scovillei*, CA1 and CA2, have been identified in Taiwan by the AVRDC–World Vegetable Center [11, 12]. CA2 is more virulent than CA1 in most tested cultivars of *Capsicum* species [9, 11]. CA2 pathotype breaks down the resistance of *Capsicum chinense* PBC932-dervied lines, which are resistant to CA1 pathotype [9, 11, 12]. CA2 distribution was limited before the year 2000, but since then has become dominant and replaced CA1 in most of the chili pepper planting locations in Taiwan [11, 12]. According to amplified fragment length polymorphism (AFLP) analysis, CA2 members are homogenous and mostly clonal, whereas CA1 are genetically diverse [11]. Strains Coll-153 and Coll-365 are grouped in CA1 pathotype and strain Coll-524 is a member of the CA2 pathotype. Understanding the variation among the three strains may provide insight about the high virulence of the CA2 pathotype.

Whole genome sequencing has had a significant impact on understanding the biology, ecology, genetics and evolution of various organisms. In fungi, comparative genomic study of genetic variations has revealed mobile pathogenicity chromosomes in *Fusarium* [13], evolutionary adaptations from a plant pathogenic to an animal pathogenic lifestyle in *Sporothix* [14], an influx of transposable elements creating a genetically flexible landscape to respond to environmental changes in *Pyrenophora tritici-repentis* [15], potential orthologs in adaptation to specific hosts or ecological niches in *Botrytis*, *Colletotrichum, Fusarium*, *Parastagonospora* and *Verticillium* [16–21], the evolution and diversity of putative pathogenicity genes in *Colletotrichum tanaceti* and other *Colletotrichum* species [22, 23], core and specific genetic requirements for fungal endoparasitism of nematodes in *Drechmeria coniospora* [24], host-specific and symptom-related virulence factors horizontally transferred from *Fusarium* to *Verticillium* [25, 26]. Genomic comparisons with multiple races within a species have also been performed recently in *Fusarium* and *Verticillium* and revealed that secondary metabolites are crucial factors for stunting symptom development in *F. fujikori* and defoliation symptom development in *V. dahliae* [26, 27]. In addition, a study on 18 isolates of *V. dahliae* identified highly variable regions with race specific signatures [28]. However, most of these studies were in-silica analyses. Only few included genetic studies to demonstrate the functions of the potential factors selected by in-silica analysis, and these studies were on *Fusarium* and *Verticillium* [18, 25–27].

In our previous study, we investigated the plant-pathogen interactions between the chili pepper fruit and the three *C. scovillei* strains Coll-153, Coll-365 and Coll-524 and revealed the infection process and several potential virulence factors determining the infection and colonization of host tissues [9]. On the original chili pepper host, Coll-524 usually shows higher virulence than the other two strains. Coll-365 shows significantly slower growth on agar medium and has the weakest virulence on *Capsicum spp.*. Coll-365 produces less spores in axenic culture and *in planta*. Coll-365 was found to accumulate less turgor pressure in appressorium but produces higher levels of cutinase activity than the other two strains. Examination of the infection process showed that the three strains can form primary hyphae in the epidermal cells at 72 h post-inoculation (hpi), indicating no delay in the penetration step for Coll-365 compared to Coll-524 and Coll-153. The three strains can grow into the cuticle layer of chili pepper fruit at 24 hpi and form a highly branched structure (HBPS) within the cuticle layer and penetrate epidermal cells at 72 hpi [29]. After infecting the epidermal cells of chili pepper fruits, Coll-365 appeared to have less ability to colonize host tissues, formed limited lesions on infected tissue and produced much fewer spores on the infected tissue. In addition, Coll-524 was more resistant to host defense compound capsaicin than Coll-153 and Coll-365. In this previous work, the genetic variations and genes contributing to the defects of growth and virulence remained undeciphered.

In this study, to decipher the genetic variations of the three *C. scovillei* strains, we sequenced the genomes of the three strains, investigated the differences in their genome compositions, identified potential genes involved in the phenotypes, and verified genes involved in the defects of growth and virulence of Coll-365. We setup a simple mathematical method to identify ORF variations and successfully identified DNA fragments involved in the defects in growth and virulence of Coll-365. Moreover, by gene transformation we have demonstrated four genes that function in germination, growth and/or virulence of the chili pepper pathogen *C. scovillei*.

## Results

### Phylogenetic analysis revealed that Coll-524, Coll-153 and Coll-365 are closely related to *C. scovillei*

To understand the phylogenetic relationships among the three *Colletotrichum* strains, multi-locus phylogenetic analysis with 174 fungal strains was performed. The results indicated that Coll-153, Coll-365 and Coll-524 all belong to the acutatum species complex **(S1 Fig)**. Further examination of the original hosts of members in acutatum species complex revealed that the three strains formed a small clade with other strains and most of the members were isolated from *Capsicum* plants (**S2 Fig**). However, the phylogenetic tree generated from the 173 *Colletotrichum* strains could not distinguish the phylogenetic relationship of the three strains with *C. scovillei* (CBS 126529, the holotype strain of *C. scovillei* [2]) and C. *guajavae* (IMI 350839, the holotype strain of *C. guajavae* [2]) **(S2 Fig)**. Another multi-locus phylogenetic analysis was conducted with 18 strains **(S1 Table)**, including 12 strains from **S2 Fig** and 6 strains (marked with #) that were closely related to *C. scovillei* CBS 126529 and *C. guajavae* IMI 350839 in the clade 2 of the *Colletotrichum acutatum* species complex [2]. The results indicated that the three strains were closer to *C. scovillei* CBS 126529 than *C. guajavae* IMI 350839 **(Fig 1A)**. In addition, Coll-524 was closer to *C. scovillei* TJNH1 isolated from pepper in China and two *C. fioriniae* strains (HC89 and HC91) isolated from apple in the USA than Coll-365 and Coll-153 **(Fig 1A)**. These results indicated that strains Coll-524, 153 and 365 all belong to the *C. scovillei* species. In addition, the three strains showed significant differences in growth and virulence but had a similar preference for temperature range (**Fig 1B-1D and** [9]).

**Figure 1.**
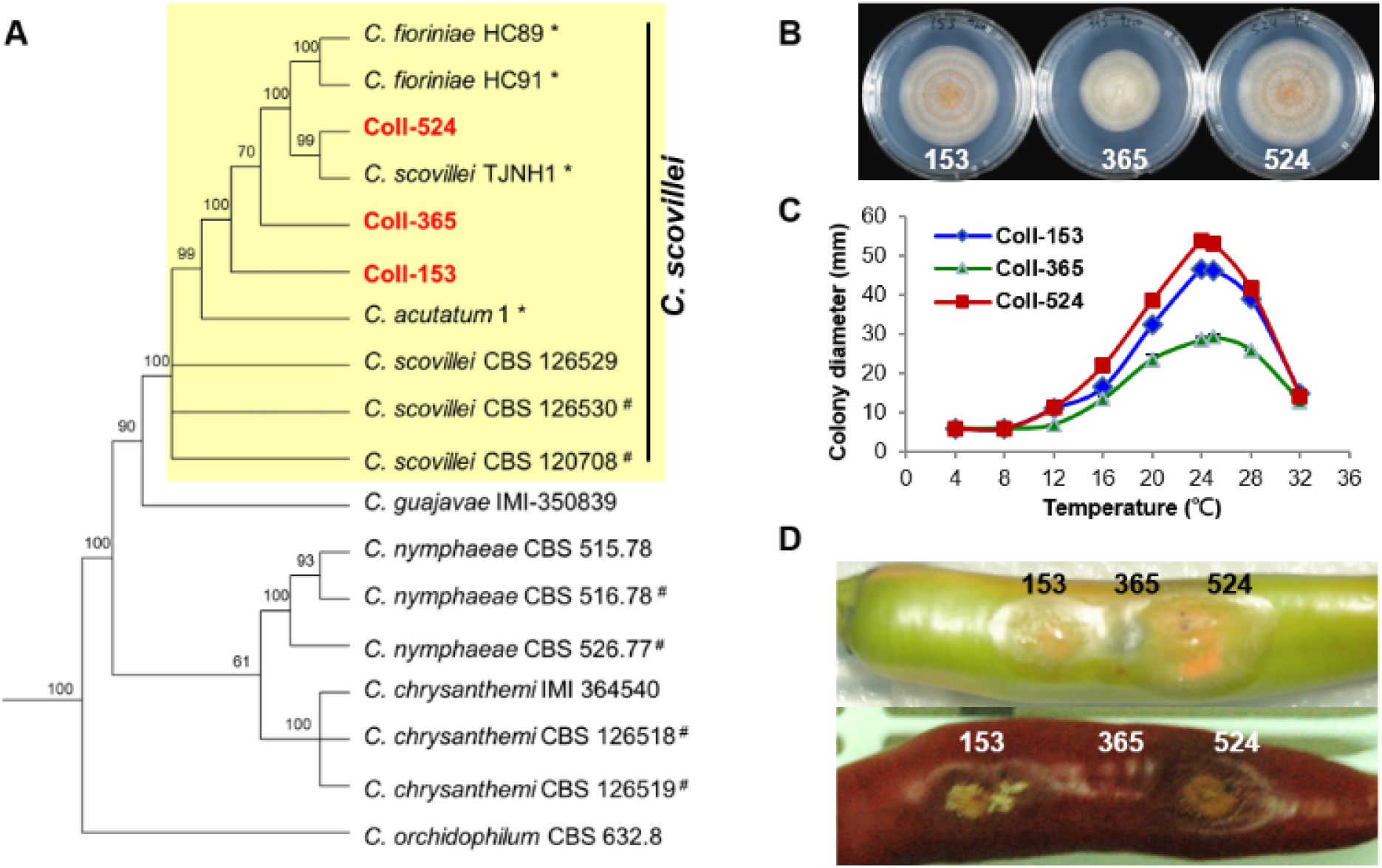
Phylogenic relationships (A), colony morphology (B), growth (C), and virulence (D) of strains Coll-524, Coll-153 and Coll-365. (A) Multi-locus phylogenetic tree of 18 *Colletotrichum* strains of acutatum species complex was constructed with five molecular markers (ACT1, CHS1, GAPDH, ITS and TUB2). Yellow square indicated that Coll-153, Coll-365 and Coll-524 were in the same clade with *C. scovillei* CBS 126529, the holotype strain of *C. scovillei*. Detail information for the 18 strains was in S1 Table. Posterior probability support values were indicated on branches. (B) Colony morphology of the three strains culturing on MS agar media for 6 days. (C) Growth of the three strains on various temperatures on PDA for 7 days. (D) Symptom of the three strains on green and red chili pepper fruits cv. Groupzest.

### Genomic comparison revealed DNA deletion patterns among different scaffolds in the three strains

The genomic sequences of Coll-524 were generated from paired-end and mate-pair libraries and assembled and annotated using the ALLPATHS-LG [30] and MAKER [31] pipelines. Genomic sequences of Coll-153 and Coll-365 were generated from paired-end libraries and mapped to the assembled Coll-524 genome (for details, see Materials and Methods). The results of assembly and annotation are summarized in **Table 1**. The genome size of Coll-524 was 51. 491 MB and consisted of 54 scaffolds with an N50 of ∼3.6 Mb. The genome size of Coll-153 was 50.114 MB and consisted of 59 scaffolds. The genome size of Coll-365 was 49.922 MB and consisted of 59 scaffolds. A total of 15,626, 15,432 and 15,387 genes were annotated in the Coll-524, Coll-153 and Coll-365 genomes, respectively **(Table 1).** BUSCO [32] analysis for genome completeness showed that 99.8% of the conserved proteins in sordariomycetes could be identified in all three strains.

**Table 1.**
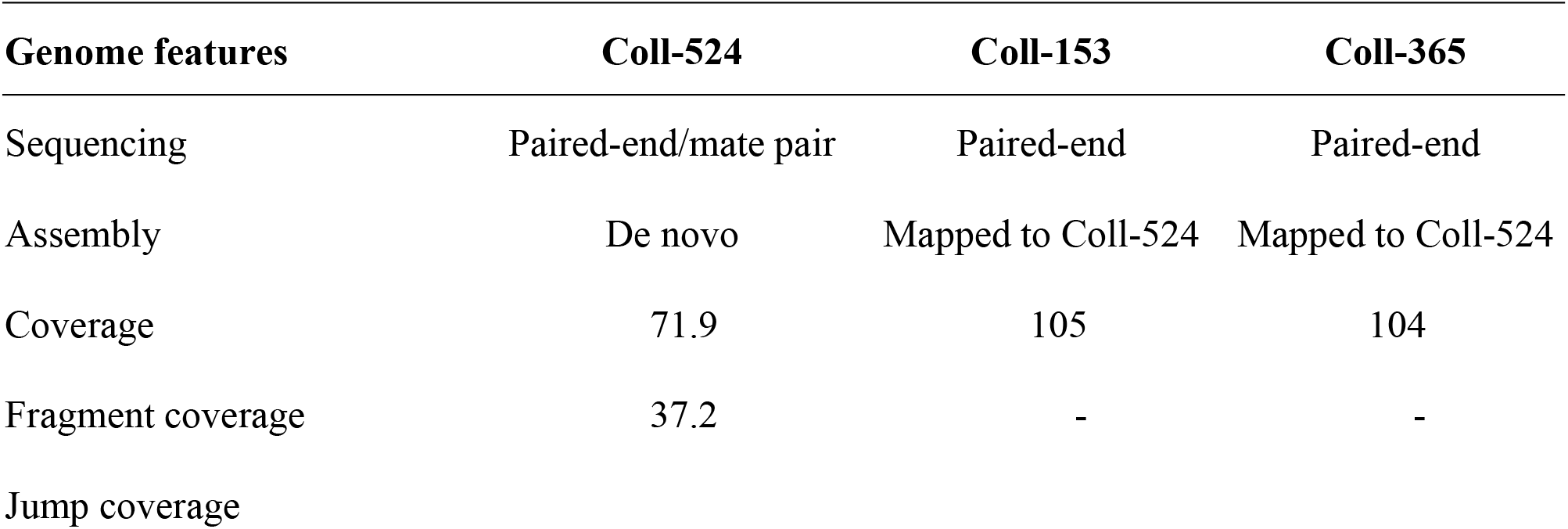
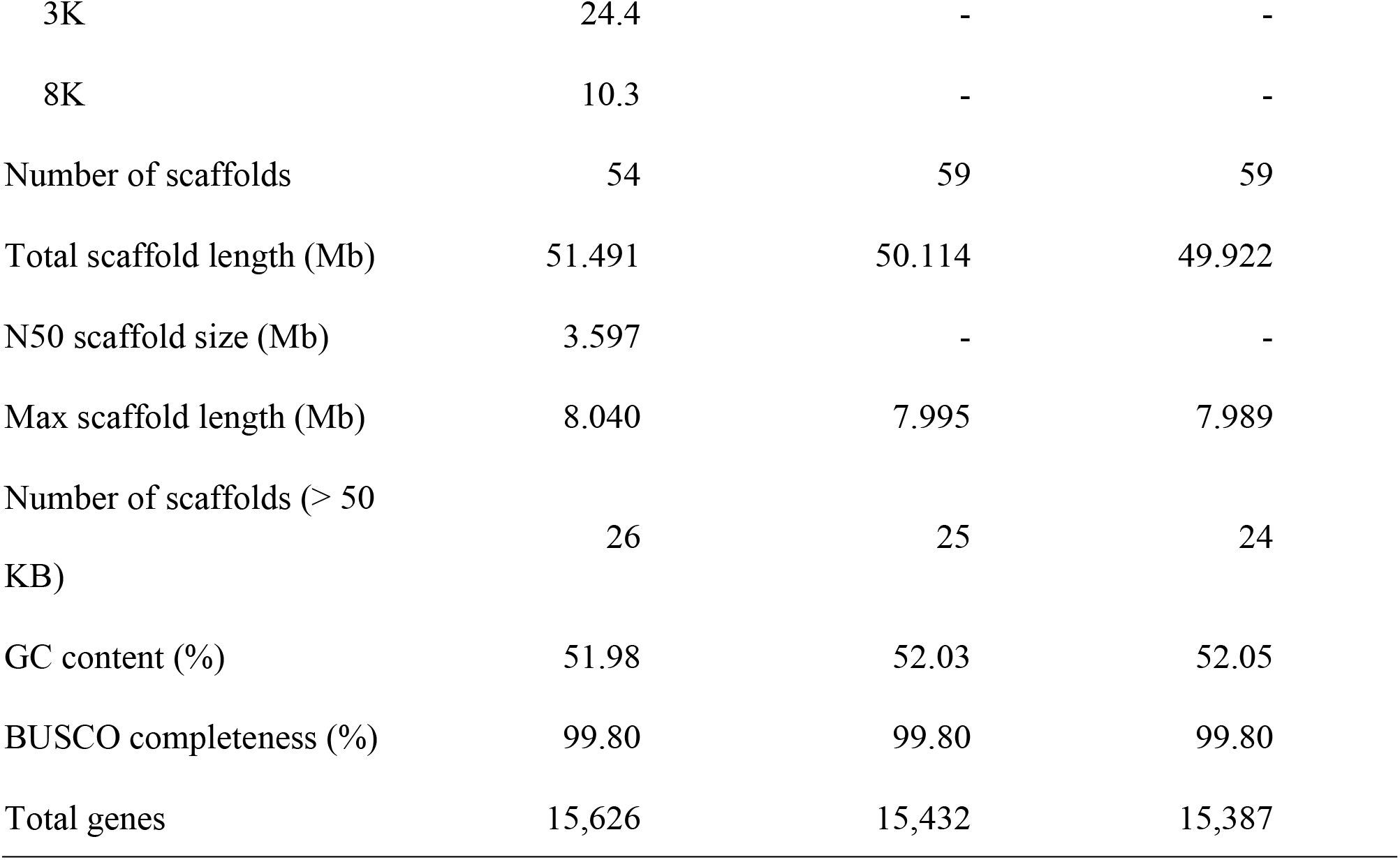
Genome features of *Colletotrichum scovillei* strains Coll-524, Coll-153 and Coll-365.

Coll-153 and Coll-365 had their own special genome fragments compared to Coll-524, and there were six scaffolds for Coll-153 and seven scaffolds for Coll-365. One (0.9-kb) and two (0.9-kb and 5.1-kb) of the 54 scaffolds in Coll-524 did not exist in Coll-153 and Coll-365, respectively, and no genes were encoded in the two scaffolds. A total of 29 and 30 genes were annotated from Coll-153 and Coll-365 specific genome sequences, respectively, and 23 of them are homologs in both Coll-153 and Coll-365. None of these genes had traits related to pathogenicity when blasting to the PHI database.

Genome mapping of Coll-153 and Coll-365 to Coll-524 was analyzed with CLC Genomic Workbench v.9.5.1 and provided the specified information of DNA insertion, deletion, removal, and miscellaneous differences (misc. differences, which included single nucleotide polymorphism (SNP) and multiple nucleotide polymorphism (MNP) in this study; see Materials and Methods) in Coll-153 and Coll-365 **(Table 2)**. The results showed that Coll-153 and Coll-365 had similar patterns within the four types of variations, but Coll-365 had a slightly higher number of total variations than Coll-153. The removed sequence database (RSD; see Materials and Methods) showed that 991,585 and 1,179,624 bp were absent in Coll-153 and Coll-365, respectively, compared to Coll-524. The largest removed fragments were 28,776 bp in Coll-365 and 10,125 bp in Coll-153. Scaffolds 19, 20 and 22 in Coll-153 and Coll-365 showed high numbers of DNA sequence removal compared to the three scaffolds in Coll-524 **(Fig 2)**. The total number of sites of DNA sequence removal in scaffolds 19, 20 and 22 were 377, 386, and 256 in Coll-153, and 167, 114 and 76 in Coll-365, respectively. A notable difference in DNA sequence removal patterns between Coll-153 and Coll-365 was that DNA sequence removal occurred significantly in scaffolds 17 in Coll-365 but not in Coll-153 (**Fig 2**). Large DNA fragment removals occurred in Coll-365 more frequently than Coll-153 (**Table 3**). However, most of them were located at the short scaffolds 19, 20 and 22. Coll-365 had four removals with DNA fragments larger than 20 kb, but none of the large fragments was found in Coll-153. Regarding the removal of DNA fragments larger than 5 kb, a total of 66 removals appeared in Coll-365 but only 8 in Coll-153. Small DNA fragment removals (< 1 kb) were found most often in Coll-153.

**Figure 2.**
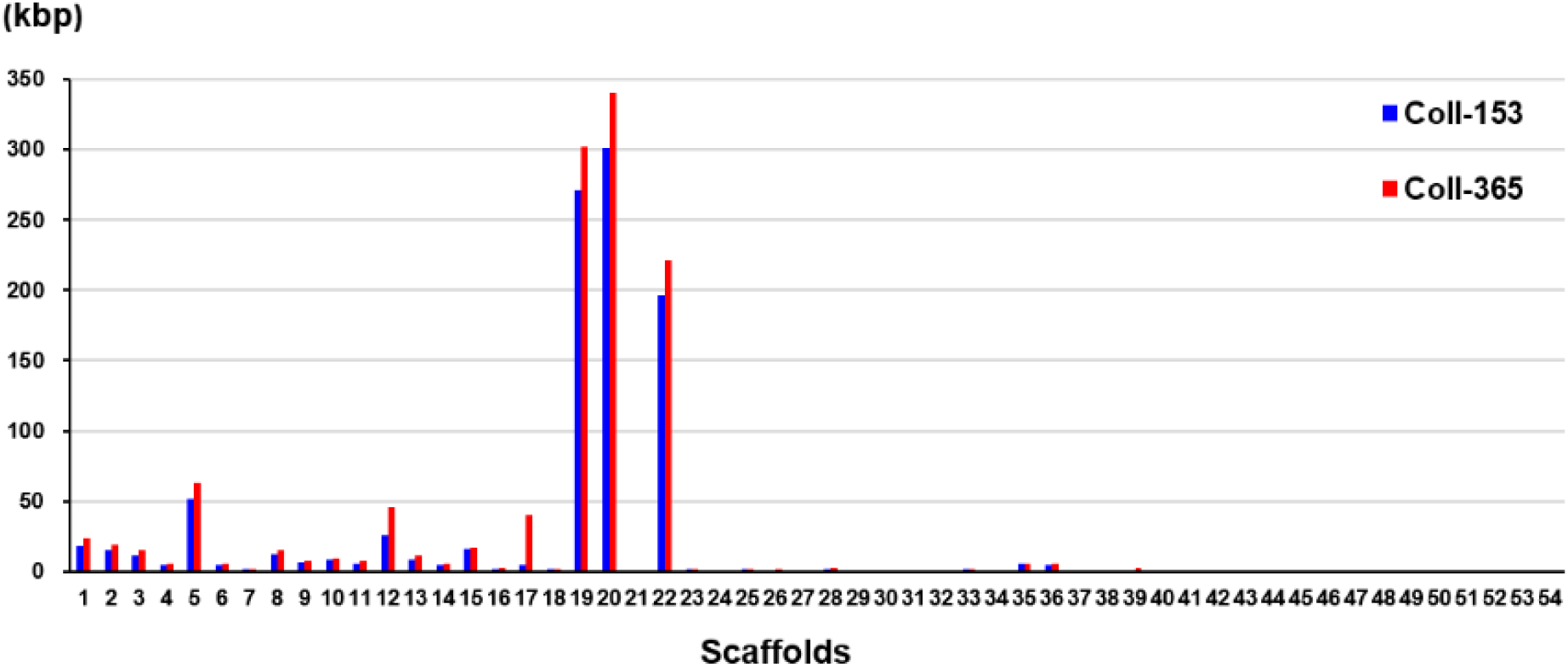
DNA sequence removal in strains Coll-153 and Coll-365 compared to strain Coll-524. Total nucleotides removed in each scaffold were indicated as blue and red bars for Coll-153 and Coll365.

**Table 2.** Genome mapping of strains Coll-153 and Coll-365 to Coll-524 analyzed with CLC to provide the datasets of DNA insertion, deletion, removed, and misc. difference in the genomes of Coll-153 and Coll-365.

| Variation Datasets | No. of variation sites or nucleotides |  |
| --- | --- | --- |
|  | Coll-153 | Coll-365 |
| Insertion |  |  |
| Insertion sites | 241 | 361 |
| Insertion bps | 338 | 506 |
| Deletion |  |  |
| Deletion sites | 302 | 451 |
| Deletion bps | 417 | 618 |
| Misc. difference<sup>a</sup> |  |  |
| Misc. difference sites | 7,411 | 9,833 |
| Misc. difference in bps | 9,179 | 12,218 |
| Removed |  |  |
| Removed sites (N<sup>b</sup><20% of the removed region) | 1,623 | 731 |
| Removed bps (N<sup>b</sup><20% of the removed region) | 991,585 | 1,179,624 |

<sup>a</sup>, the dataset of Misc. difference consisted of single nucleotide polymorphism (SNP) and polymorphism with more than one nucleotide which was named as multiple nucleotide polymorphism (MNP) in this study.
<sup>b</sup>, N indicates gaps in scaffolds.

**Table 3.** Large DNA fragment removal statistic of strains Coll-153 and Coll-365.

| DNA removal | Coll-153 | Coll-365 |
| --- | --- | --- |
| $\geq 20$ kb | 0 | 4 |
| $\geq 10$ kb | 1 | 21 |
| $\geq 5$ kb | 7 | 41 |
| $\geq 1$ kb | 325 | 175 |
| <1 kb | 1290 | 490 |
| Sum | 1623 | 731 |

### ORF variation analysis identified ORF losses in strains Coll-153 and 365

DNA insertion, deletion, removal, and misc. differences (SNP and MNP) occurring in Coll-153 or Coll-365 may affect the open reading frames (ORFs) of their genes. To understand the influence on ORFs, we used a simple mathematical method to detect ORF variation (ORF-V) for all ORFs in Coll-153 and Coll-365 as illustrated in **Fig 3** and described in Materials and Methods. The results showed that most of the ORF-Vs occurred with SNP in Coll-153 and 365, and Coll-365 appeared to have significantly greater numbers of full ORF losses than Coll-153 **(Table 4)**. The total numbers of ORF-V of Coll-153 and Coll-365 compared to Coll-524 were 708 and 725, respectively, and 799 of them were singular. A total of 91 and 74 ORF-Vs were solely found in Coll-365 and Coll-153, respectively, but most of these ORF-Vs were SNP in Coll-153 or Coll-365, indicating that the two strains had similar genes with ORF-V to Coll-524. The major difference in the two strains with regard to ORF-V was found in scaffold 17 and was caused by ORF deletion as described below. The 799 singular ORFs were used for further analysis.

**Figure 3.**
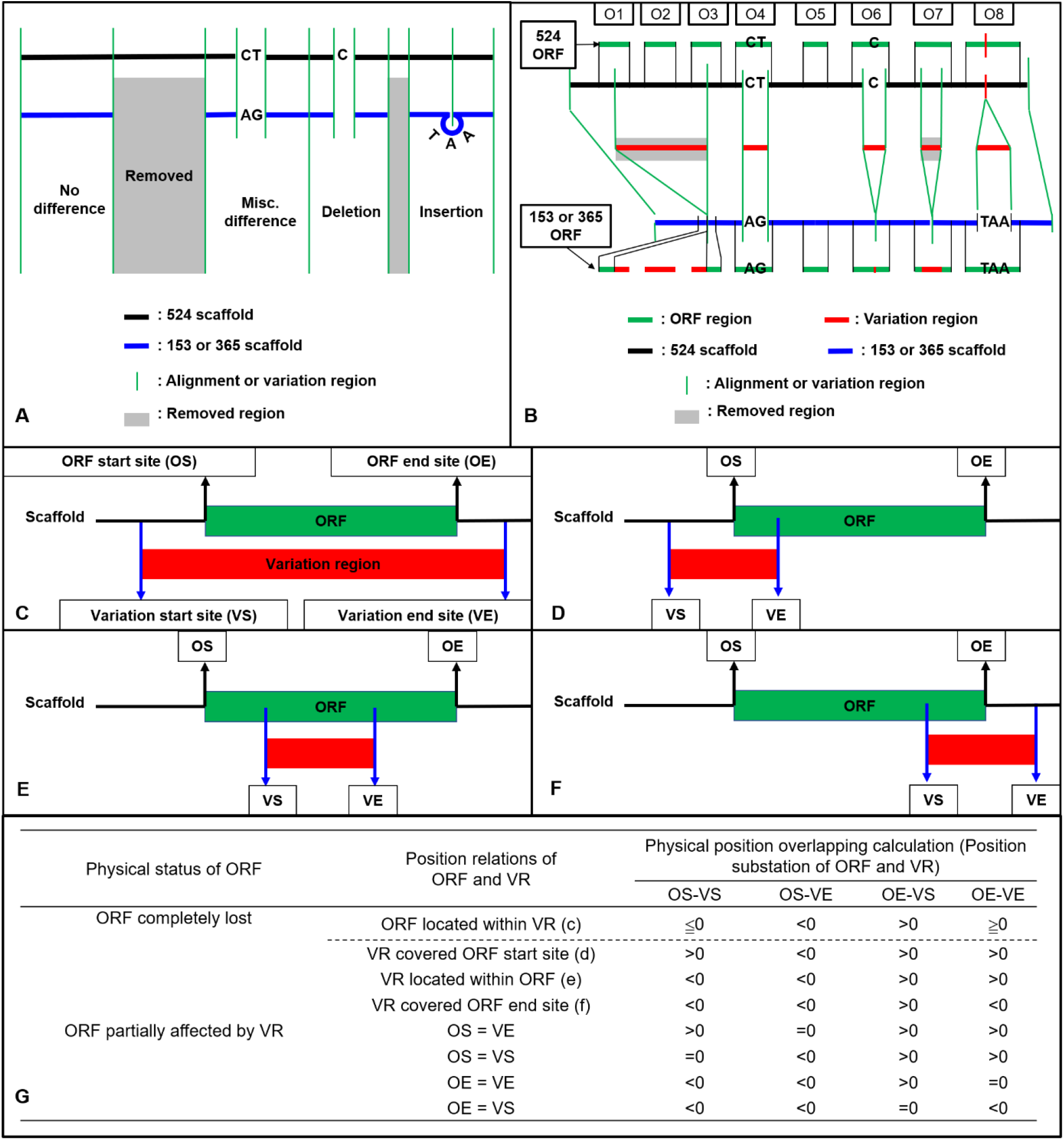
Strategy illustration (A-B) and calculation method (C-G) used for ORF variation identification. (A) Variations were generated after reads mapping of Coll-153 or Coll-365 to Coll-524. The mapping could result in four types of variations (Removed, Misc. Difference, Deletion and Insertion). (B) ORF regions of Coll-524 were compared to variation regions to find the overlappings. An ORF with the overlapped region indicated that this ORF is varied between the two strains. (C-F) Calculation methods used to identified all potential position relations between an ORF and variation regions. (G) Calculation for position difference to identify ORF variation types, removed or partial differences. VR, variation region; OS, ORF start site; OE, ORF end site; VS, variation start site; VE, variation end site.

**Table 4.**
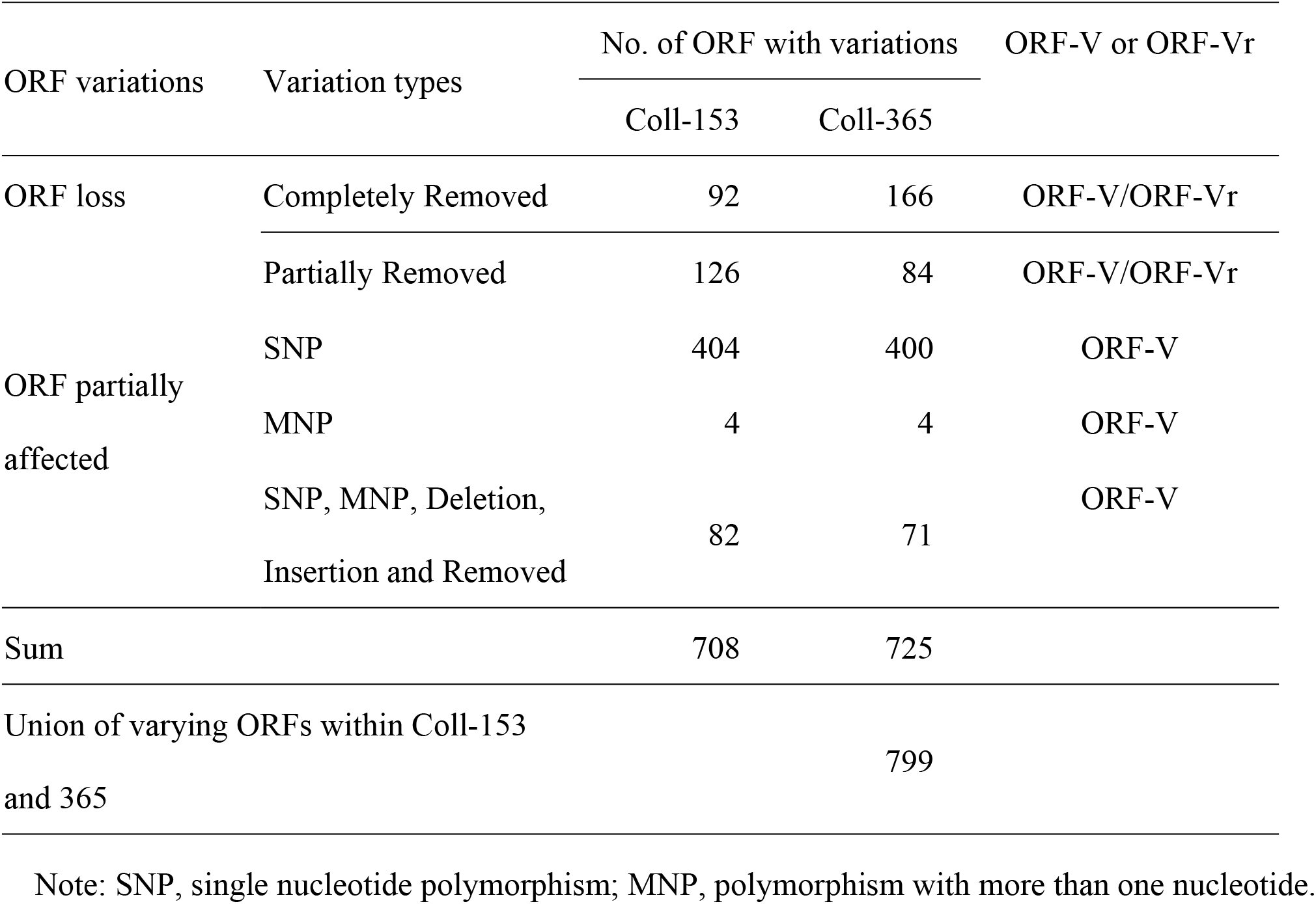
ORF variation statistics of strains Coll-153 and Coll-365 detected and calculated based on the method shown in Fig 4.

ORFs affected by DNA removal (ORF-Vr) in each scaffold were further analyzed and the results are displayed in **Fig 4**. ORF-Vr mostly occurred in scaffolds 19, 20 and 22 in Coll-153 and 365. There were a total of 260 and 274 ORF-Vr events in Coll-153 and Coll-365, respectively. All ORFs were ORF-Vr in scaffolds 19, 20, 22 and 44; however, scaffold 44 was a short scaffold with only one ORF (**Fig 3**). The distribution of ORF-Vr in Coll-153 and 365 was very similar except that 14 ORFs clustered in a 34-kb genomic fragment were totally or partially removed in Coll-365 compared to 1 ORF with SNP only in Coll-153 in scaffold 17 (**Fig 5**). The 14 ORFs encode 4 transcription regulation-related proteins, 1 GPI-anchored protein, 4 enzymes and enzyme-related proteins, and 5 hypothetical proteins. In addition, when compared with 62 genomes of other *Colletotrichum* strains that are available in the NCBI database, the 14 genes appeared in almost all members of the acutatum species complex, except Coll-365 (**Fig 6**). Moreover, genes 15003 and 15019 existed in nearly all assayed strains, indicating that the two genes are core genes of the *Colletotrichum* species. Five of these genes existed in almost all the strains in the acutatum species complex, suggesting the possibility of them being acutatum species complex-specific genes (**Fig 6**).

**Figure 4.**
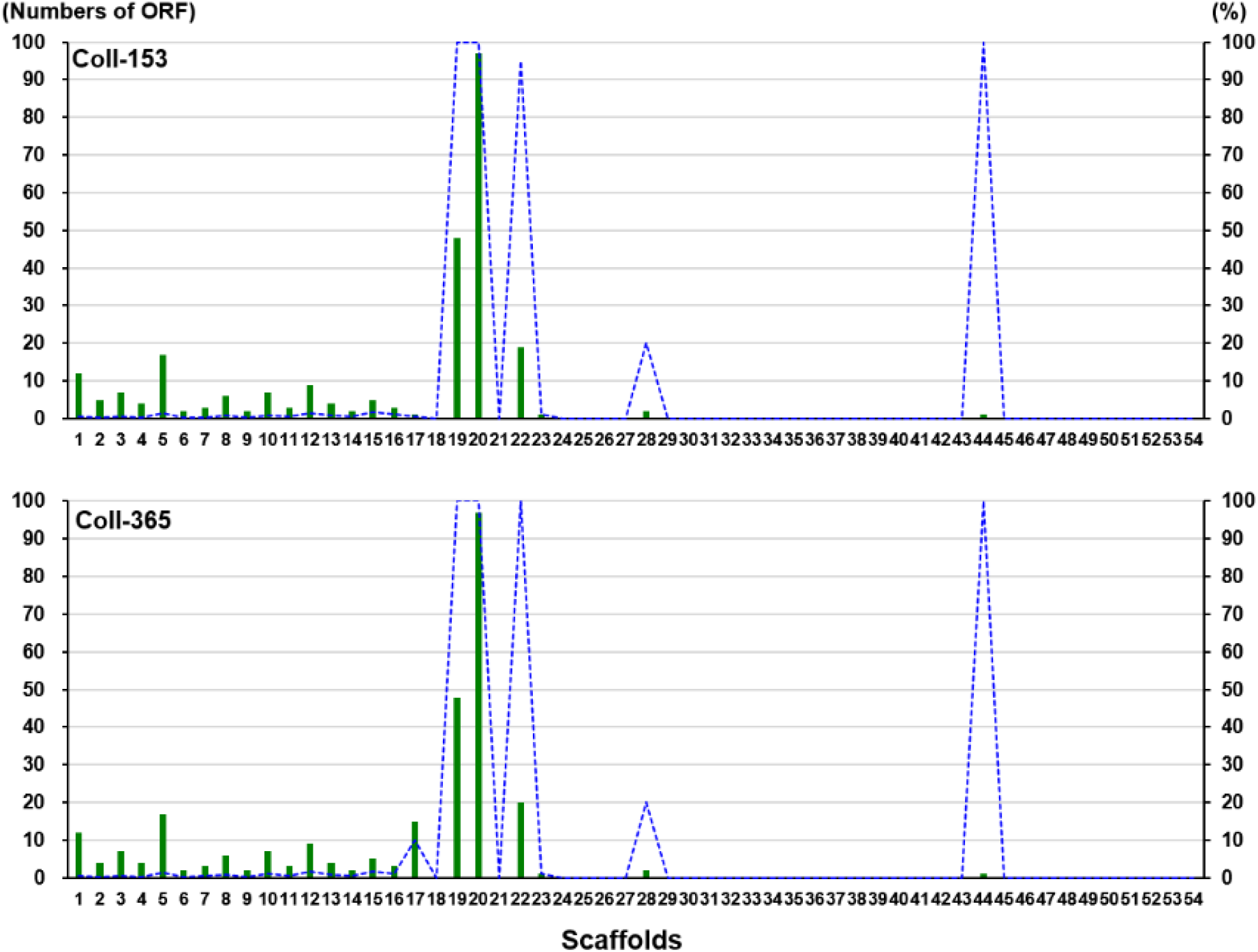
ORF-Vr in strains Coll-153 and Coll-365 compared to strain Coll-524 in each scaffold. Total ORF-Vr in each scaffold were indicated as green bars. The percentage of ORF-Vr in each scaffold by comparing with ORFs of Coll-524 in each scaffold was indicated with blue lines.

**Figure 5.**
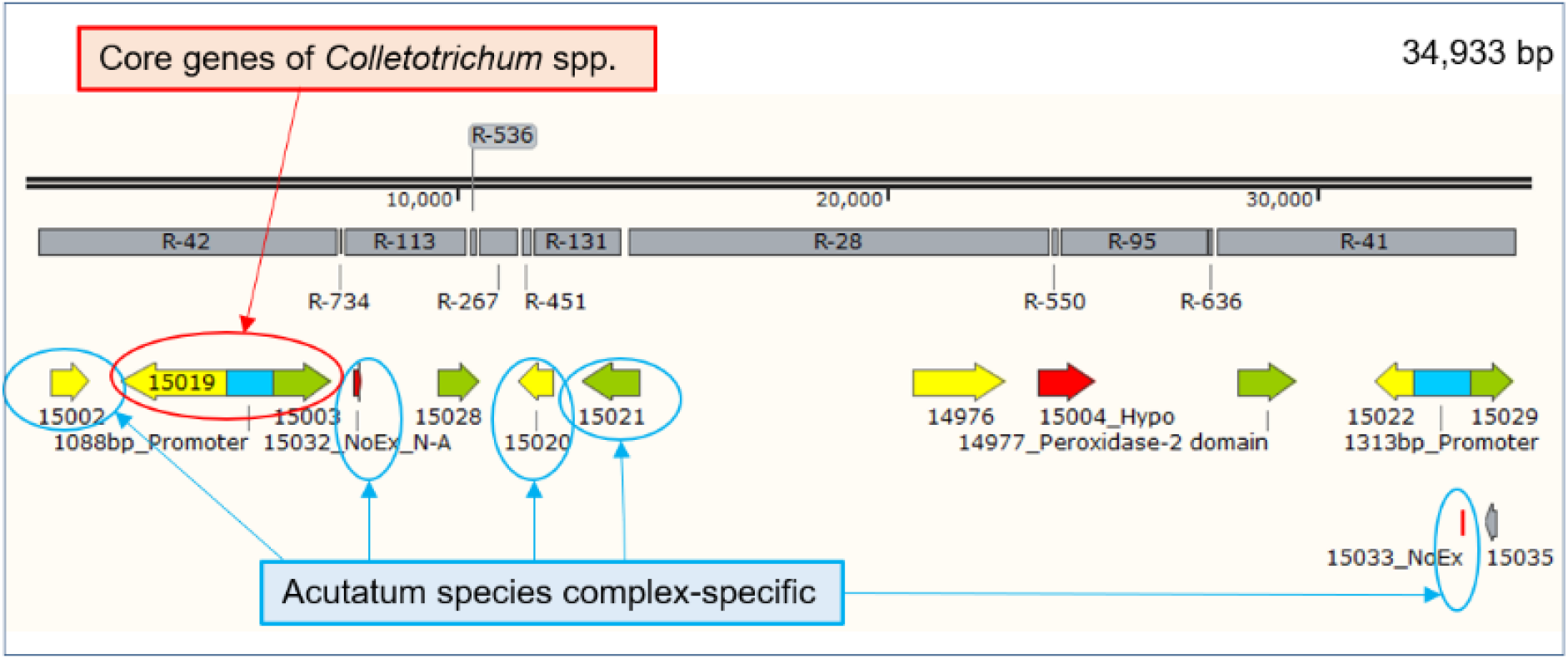
Genetic map of the 14 removed genes in scaffold 17 of strain Coll-365. The gray bars with R-number indicated the removed fragments in Coll-365 compared to Coll-524. The 14 genes located in the 34.9-kb fragment of scaffold 17 are genes 15002, 15019, 15003, 15032, 15028, 15020, 15021, 14976, 15004, 14977, 15022, 15029, 15033 and 15035. NoEx: Expression data are non-available.

**Figure 6.**
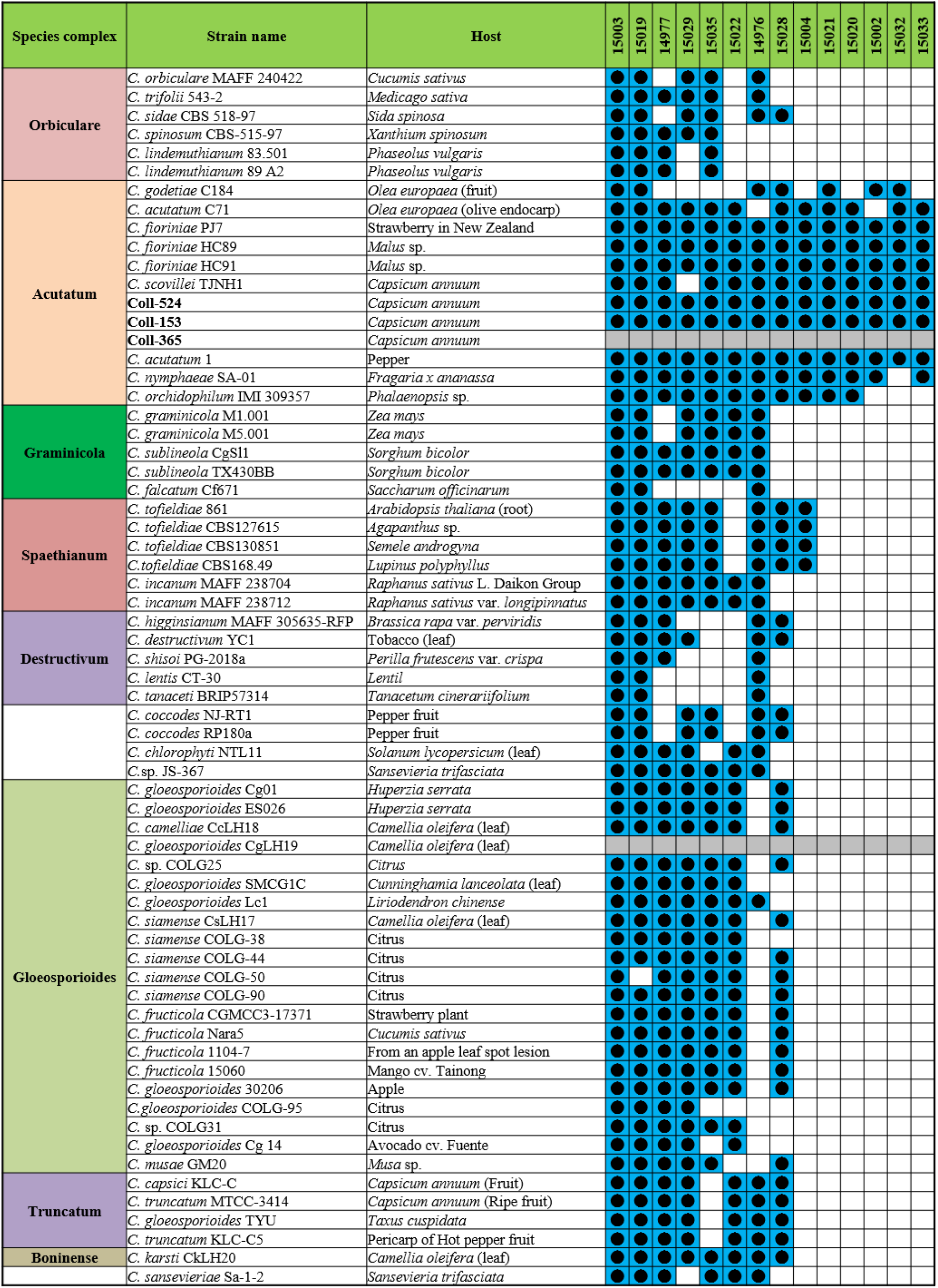
The distribution of the 14 genes of scaffold 17 in strains Coll-524, 153, and 365, and 62 *Colletotrichum* strains. The 14 genes were analyzed using nucleotide BLAST against the 62 genomes of *Colletotrichum* strains. Blue squares indicate the presence of genes by the blasting with coverage >90% and identity >60%. The gene was ordered from left to right by distribution frequency in the 62 genomes.

### Gene ortholog analysis showed that the three strains carry slightly different ortholog compositions

To understand the difference in orthology among the three strains, proteins were analyzed using OrthoFinder [33]. Ortholog analysis showed a high similarity in gene compositions among the three strains as shown in **Table 5**. More than 88% genes in the three strains belonged to single-copy orthogroups, indicating that the three strains all carried the 13,779 single-copy genes (**Table 5**). A total of 502 multiple-copy orthogroups with the same gene numbers containing 1,167 genes were all identified in the three strains. A combination of single-copy orthogroups and multi-copy orthogroups with the same gene copy number showed that there were a total of 14,946 genes in the three strains and they were 95.7, 96.9 and 97.1% of the total genes in Coll-524, 153 and 365, respectively. This suggests that the three strains had high similarity in gene compositions **(Table 5)**. The ortholog differences within the three strains analyzed using the dataset of multi-copy orthogroups with different gene number showed Coll-524 carried considerably greater numbers of genes than the other two strains. The three strains all carried the 128 orthogroups but they had different gene numbers in each of the orthogroups. Among the 128 orthogroups, Coll-524 had 97 and 101 more genes than Coll-153 and Coll-365, respectively, and there were a total of 103 singular genes found from the 97 and 101 genes. Regarding the differences in orthogroups that only existed in two strains, 92, 33 and 94 orthogroups were identified in Coll-524 and Coll-153, Coll524 and Coll-365, and Coll-153 and Coll-365, respectively (**Table 5**). Further analysis of the differences between pairs of two strains showed that some orthogroups were only found in two strains, and Coll-524 and Coll-153 shared more orthogroups (92 orthogroups, 94 genes) than Coll-524 and Coll-365 (33 orthogroups, 33 genes). Based on the data mentioned above, the combinations of the 103 genes, the 94 and 33 genes, the 85 of strain-specific genes and the 196 unassigned genes of Coll-524 strain, 511 genes in total, were used together with other criteria for further gene selections used in gene functional analysis.

**Table 5.**
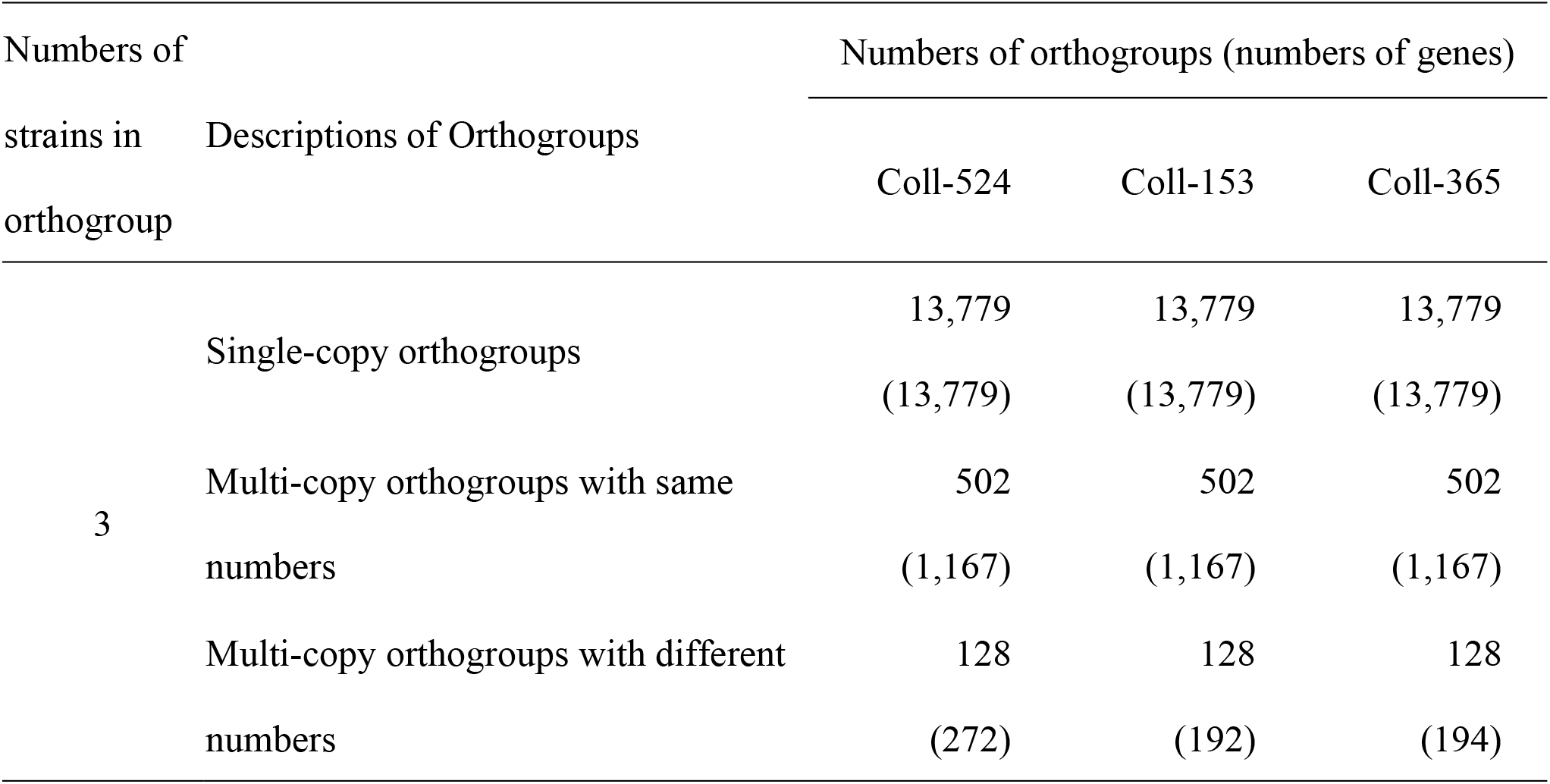
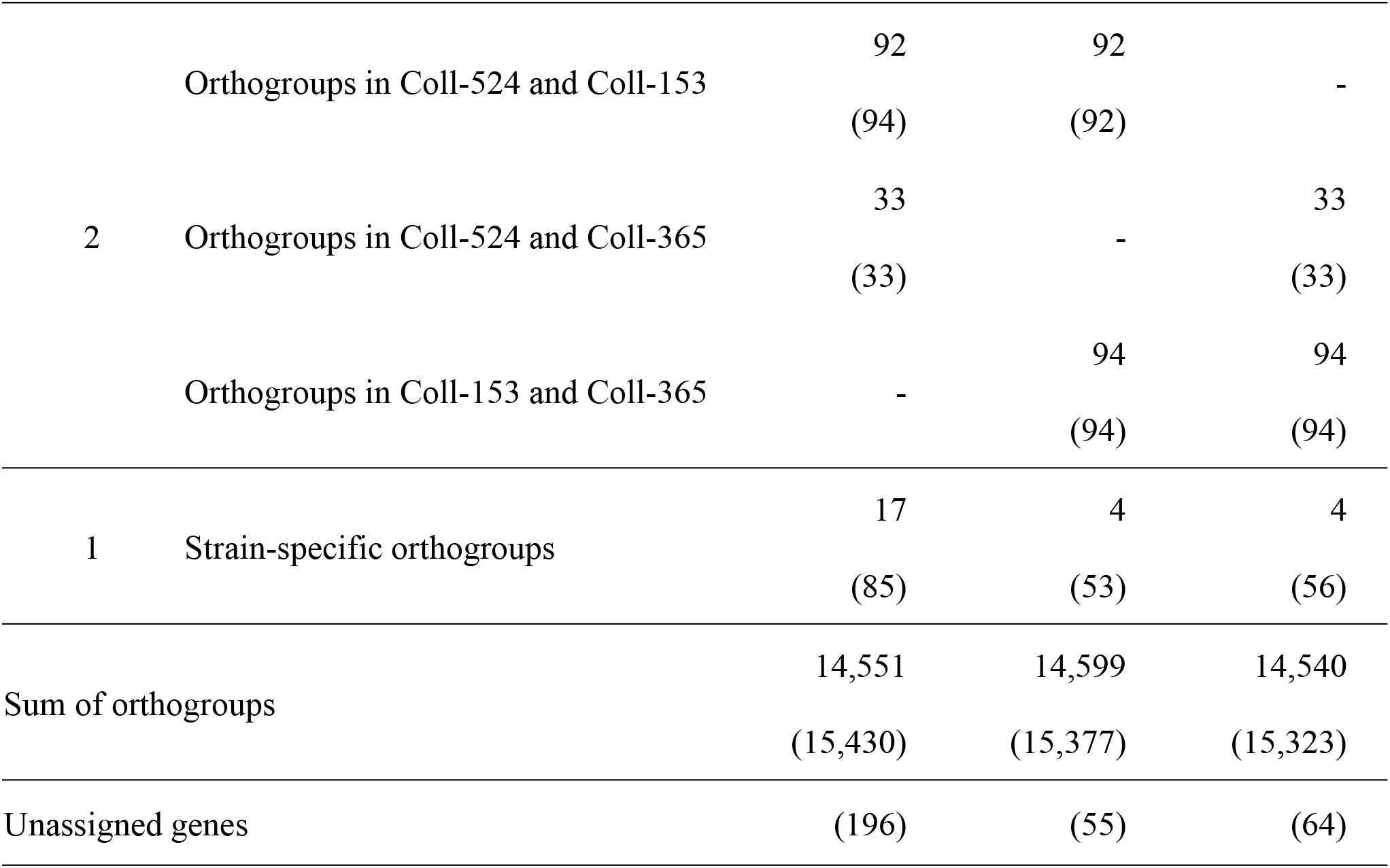
Orthogroup types and distributions of *Colletotrichum scovillei* strains Coll-524, Coll-153 and Coll-365.

### Comparison analysis revealed variations in pathogenicity-related categories in the three strains

To understand the variations in pathogenicity-related genes among the three strains, analyses of pathogenicity-related functional categories were performed. Six categories were used, effectors, carbohydrate active enzymes (CAZymes), secondary metabolism clusters, transcription factor (TF), and enzymes in the Kyoto Encyclopedia of Genes and Genomes (KEGG) pathway, and pathogen-host interaction (PHI) related genes. The results showed the three strains had very similar numbers of genes in various categories **(S2 Table)**. When combining these data and the ortholog data in a further analysis, notable differences were observed **(Table 6 and S3 Table)**. With regard to strain-specific orthogroups and unassigned genes in ortholog analysis, Coll-524 had significantly greater gene numbers of effectors and PHI than Coll-153 and 365 **(Table 6)**. With regard to multi-copy orthogroups with different gene numbers, Coll-524 had more genes belonging to the six functional categories than Coll-153 and 365 **(S3 Table)**. For orthologs only found in two strains, there were no notable differences between any set of two-strains with regard to gene numbers among the six functional categories.

**Table 6.**
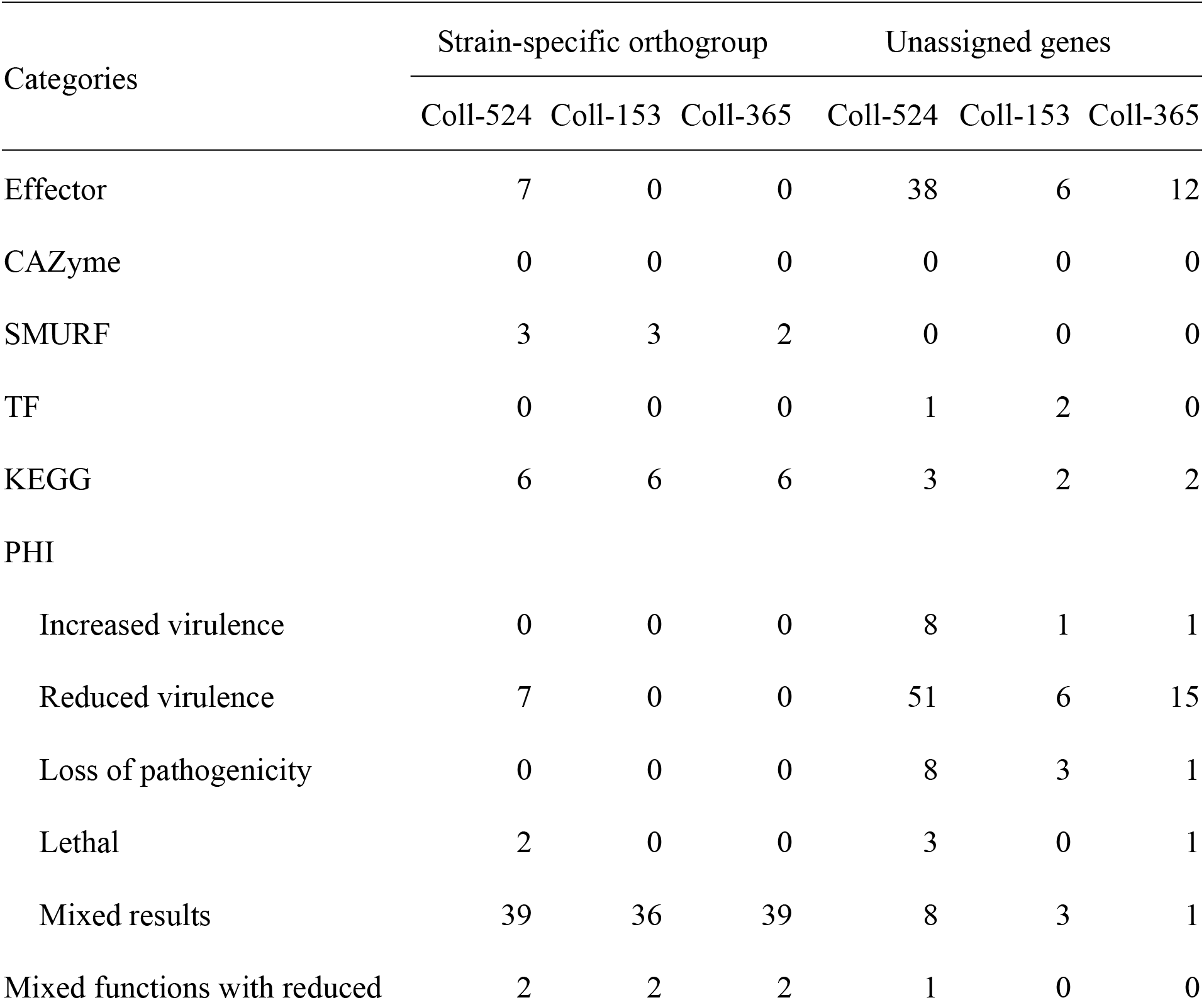
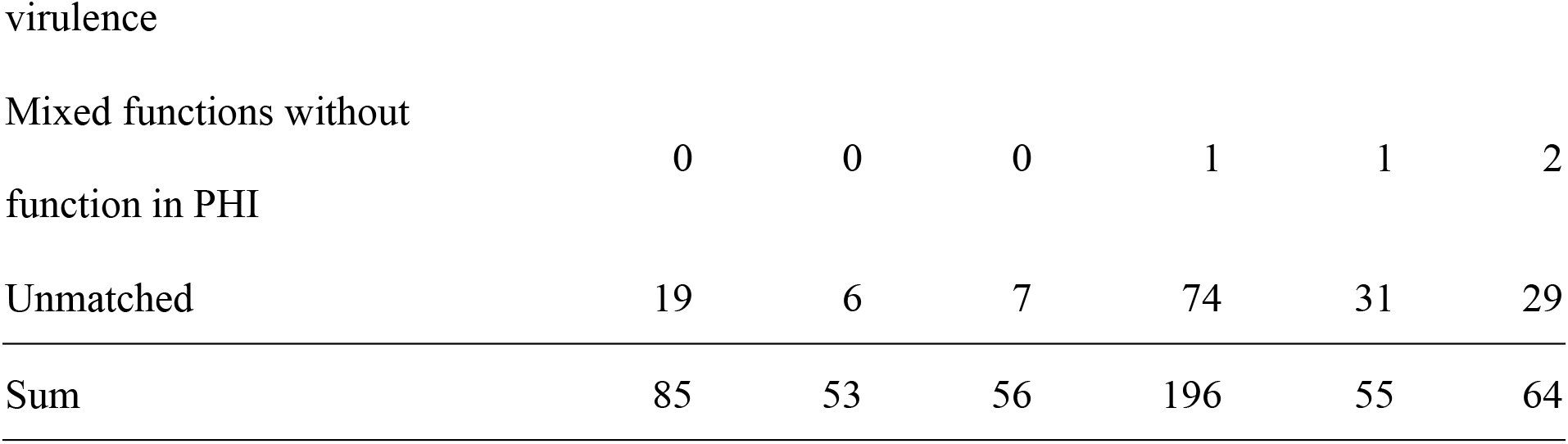
Functional category statistics of strain-specific orthogroups and unassigned genes of the three *Colletotrichum scovillei* genomes.

The gene variations of the six functional categories in scaffolds 17, 19, 20 and 22 in the three strains are summarized in **Table 7**. Coll-153 and Coll-365 had remarkably fewer genes than Coll-524 in PHI (114 genes) and effector (33 genes) categories (**Table 7**). The ORFs of the 33 effector genes were further analyzed using nucleotide BLAST (blastn) against 62 genomes of *Colletotrichum* strains with genomes available in the NCBI database. As shown in **Fig 7**, two strains of *C. acutatum* species complex (Coll-524 and *C. scovillei* TJNH1) carried all the 33 effector genes, while one strain of this complex, *C. acutatum* 1, had 32 of the effectors. However, two strains with phylogenetics closely related to Coll-524, *C. fioriniae* HC89 and HC91, only carried two of the 33 effector genes. Among the 62 *Colletotrichum* strains, 27 did not carry any of the 33 effectors and they were mainly in the species complexes graminicola, spaethianum, and gloeosporioides. Interestingly, none of the members of the graminicola species complex carried any of the 33 effectors. Moreover, a total of 24 of the 33 effectors were only found in the acutatum complex, and not in the other complexes.

**Figure 7.**
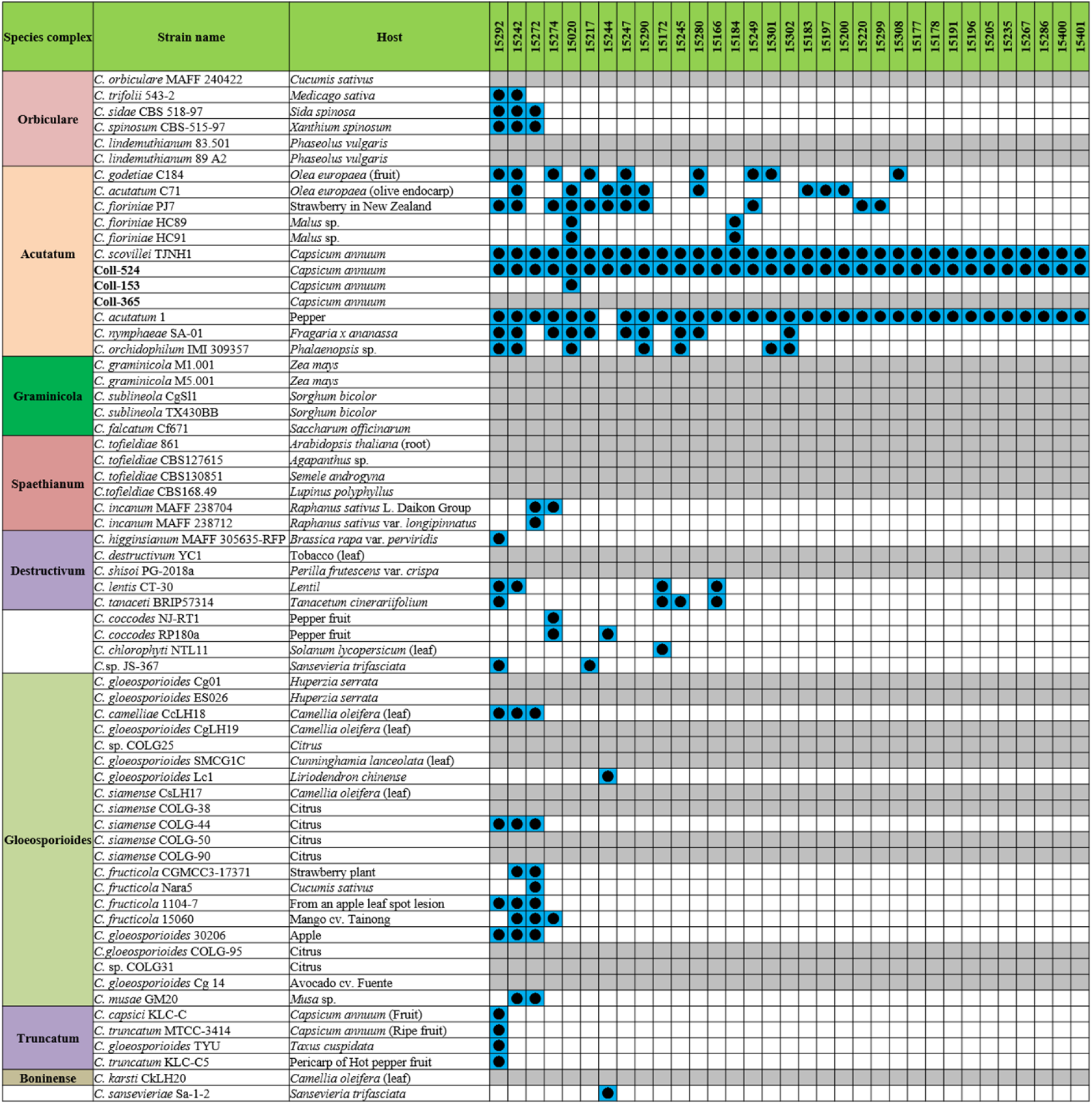
The distribution of the 33 effector genes in strains Coll-524, 153, and 365, and 62 *Colletotrichum* strains. The 33 effector genes were analyzed using nucleotide BLAST against the 62 genomes of *Colletotrichum* strains. Blue squares indicate the presence of genes by the blasting with coverage >90% and identity >60%. The gene was ordered from left to right by distribution frequency in the 62 genomes.

**Table 7.** Functional category statistics of genes located at scaffolds 17, 19, 20 and 22 of *Colletotrichum scovillei* strains Coll-524, Coll-153 and Coll-365.

| Categories | Coll-524/Coll-153/Coll365 |  |  |  | Removed in Coll-153<br>and /or Coll-365 |
| --- | --- | --- | --- | --- | --- |
|  | S17 | S19 | S20 | S22 | S17+S19+S20+S22 |
| Effector | 20/19/19 | 11/0/0 | 19/0/1 | 2/0/0 | 33 |
| CAZyme | 8/8/7 | 0/0/0 | 0/0/0 | 0/0/0 | 1 |
| SMURF | 0/0/0 | 7/0/0 | 0/0/0 | 0/0/0 | 7 |
| TF | 6/4/4 | 1/0/0 | 3/0/0 | 1/1/0 | 7 |
| KEGG | 10/10/9 | 2/1/0 | 4/0/0 | 0/0/0 | 7 |
| PHI |  |  |  |  |  |
| Increased virulence | 7/7/7 | 4/1/0 | 3/0/0 | 1/0/0 | 8 |
| Reduced virulence | 61/61/54 | 21/0/0 | 42/3/2 | 11/0/0 | 79 |
| Loss of pathogenicity | 8/7/6 | 6/0/0 | 5/1/0 | 0/0/0 | 12 |
| Lethal | 4/4/4 | 2/0/0 | 3/0/0 | 1/0/0 | 6 |
| Mixed results | 8/8/8 | 2/0/0 | 6/0/0 | 1/0/0 | 9 |

The variations of PHI genes among the three strains in scaffolds 17, 19, 20 and 22 were found to mainly occur in scaffolds 19, 20 and 22, especially genes required for fungal full virulence in the PHI database. A total of 79 genes with functions related to reducing virulence were found to be partially or fully removed in Coll-365 and 71 of them were not in Coll-153 (**Table 7**).

The variations of the three strains in KEGG pathways on scaffolds 17, 19, 20 and 22 were found to mainly occur on scaffold 20, in which 100% of the KEGG genes were lost in Coll-153 and Coll-365. The investigation of KEGG combined with ortholog analysis showed that gene number variations within the three strains in scaffolds 17, 19, 20 and 22 were found in five metabolic pathways. These pathways were metabolisms of drugs, purine, thiamine, phenylpropanoid, and N-glycan biosynthesis. Coll-524 carried more genes than Coll-153 and/or Coll-365 in the five pathways **(S3-7 Figs)**.

### Fot1 (TcMar-Fot1)-like transposon significantly decreased in strain Coll-365

Repeat sequences were analyzed with MIcroSAtellite (MISA) [34], TransposonPSI and RepeatModeler [35]. Repeat sequence analysis with MISA showed the microsatellite compositions were very similar in the three strains (**S4 Table**). TransposonPSI analysis revealed that 11 transposon families were found in this species but the three strains carried different amounts of the transposon families. Coll-524 contained all 11 families, but Coll-153 and Coll-365 only carried 9 families. In addition, Coll-365 had significantly less DDE_1 domain-containing transposon than Coll-153 and Coll-524 (**Fig 8**). Repeat sequence analysis with RepeatModeler showed that Coll-524, Coll-153 and Coll-365 had 53, 67, and 44 repeat families, respectively. The DDE_1 domain identified by TransposonPSI is from the TcMar-Fot1 family in the RepeatModeler database. The complete Coll-Fot1 sequence was identified by combining sequences provided by the TransposonPSI and RepeatModeler (**S8 Fig**).

**Figure 8.**
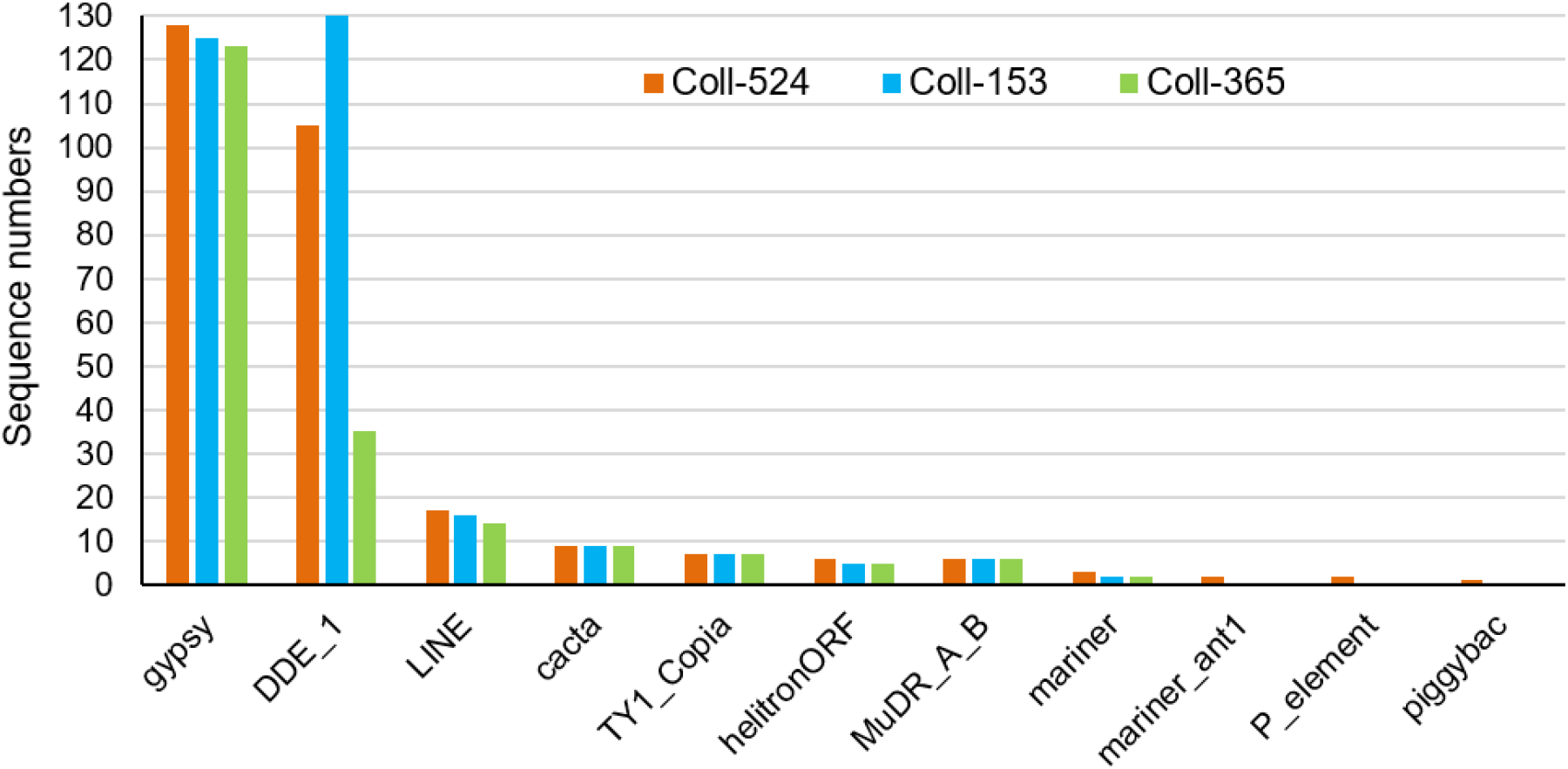
Distribution of repeat sequences among the three *Colletotrichum scovillei* strains.

### Eight genes were selected for functional analysis

Orthogroup and ORF-V analysis revealed two sets of variations in gene composition of the three strains. To obtain comprehensive analysis of variation, additional analysis was conducted by blastn using Coll-524 genes against the genome sequences of Coll-153 and 365. A total of 219 genes from Coll-153 and/or 365 with less than 50% coverage to Coll-524 were identified and used with genes selected from the other three groups, orthogroup analysis (511 genes), ORF-V analysis (799 genes) and pathogenicity-related categories analysis, for further analysis (**Fig 9**).

**Figure 9.**
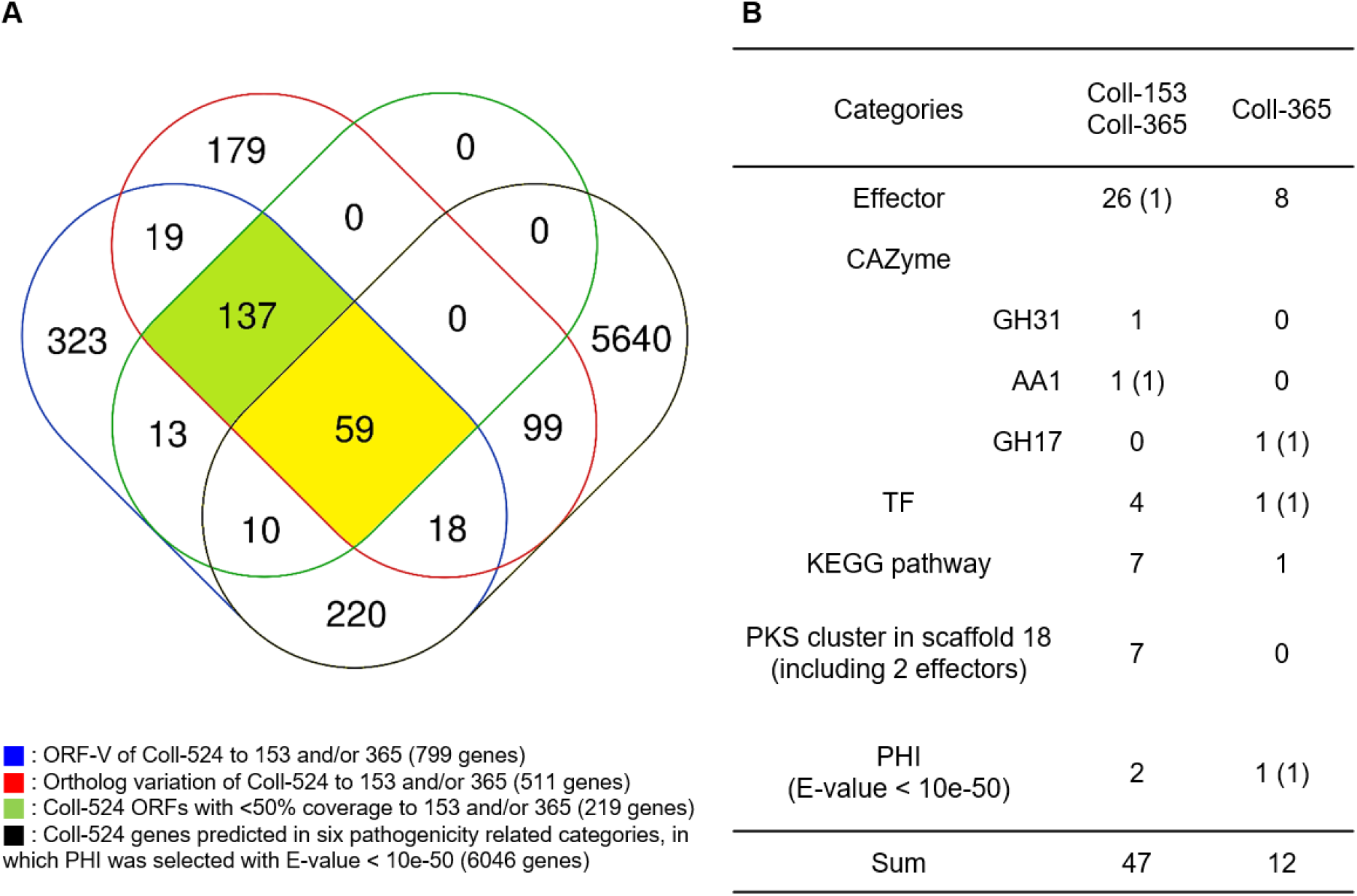
Venn diagram of selected gene groups (A) and the clustered 59 genes distribution in six function categories (B). The four selected gene groups were indicated in the bottom of panel A. In panel B, the numbers of genes disappeared in both Coll-153 and Coll-365, and in Coll-365 only were presented. The number in parentheses indicates gene numbers selected for gene functional transformation assay.

The results are shown in **Fig 9A**. A total of 59 genes were found from the four groups. Among the 59 genes, 47 genes were lost in both Coll-153 and Coll-365, and 12 genes were only absent in Coll-365, including 8 effectors, one CAZyme, one TF, one KEGG pathway, and one PHI **(Fig 9B)**. Most of the 47 genes were located at scaffolds 19 and 20, while most of the 12 genes were located at the scaffolds 17 and 20 (**S5 Table**). A total of 137 genes were clustered from three gene groups excluding the pathogenicity-related function categories (**Fig 9A**). These 137 genes contained 5 cytochrome P450 genes, 5 FAD binding domain-containing protein genes, one LysM domain-containing protein gene (15215) as well as large numbers of unknown and hypothetical protein genes. Eight genes, 5 from the 59 gene group and 3 from the 137 gene group, which were highly expressed during infection, based on RNAseq data, were then selected for functional analyses. Five genes from the 59 gene group are 11975, 14976, 14591, 15022, and 15019 (**Fig 9B**), and the 3 genes from the 137 gene group are 15003, 15029 (**Fig 5**) and the LysM domain-containing protein gene (15215). Genes encoding cytochrome P450 or FAD binding domain-containing protein were not selected because they belong to large gene families in Coll-524. The eight genes were analyzed by PCR to confirm their absence in the genome of Coll-153 and/or Coll-365 (**S9A Fig**).

### The genes lost in a 34-kb fragment have a large effect on strain Coll-365 morphology and pathogenicity

The eight selected genes were driven by their native promoters and transferred into Coll-365. Five genes were located within a 34-kb fragment at scaffold 17 and two sets of two genes were closely linked (15029 and 15022; 15003 and 15019). Therefore, for Coll-365 transgenic strains carrying two closely linked genes, three or four of the four genes were generated (**Table 8**). All transgenic strains were selected with PCR assays for the insertion fragment and/or further confirmed with RT-PCR for their expression in Coll-365 (**S9B-C Fig**). The single spore-purified transgenic strains were used for assays on spore germination, appressorium formation, growth, and pathogenicity assay on chili pepper fruits. Two independent transgenic strains of every gene transformation were used in all assays. Coll-524, Coll-153 and Coll-365 showed similar ability with regard to spore germination and appressorium formation (**Fig 10A**). Two independent transgenic strains of transformation IV (gene 15215, encoding a LysM domain containing protein; **Table 8**) showed a significantly lower germination rate than the wild-type Coll-365 in two independent experiments **(Fig 10A**). There were no significant differences in appressorium formation between the gene transformation strains and wild-type strain Coll-365. In the growth assay, Coll-365 had significantly slower growth than Coll-153 and Coll-524. Among the transgenic strains, two independent transgenic strains of transformation VIII (gene 15019), IX (carrying genes 15022, 15029 and 15019) and X (carrying genes 15022, 15029, 15019 and 15003) showed significantly enhanced growth on MS agar medium in comparison with their wild-type strain Coll-365, but still showed slightly slower growth than Coll-524 (**Fig 10B**). For pathogenicity, in a preliminary assay on chili pepper *Capsicum annuum* cv. Hero, Coll-365 was compared with transgenic strains carrying gene encoding effector, laccase, end-beta-1,3-glucasnase or LysM containing protein, and Coll-153 or Coll-524. Lesion size calculation showed that Coll-365 had significantly lower virulence than Coll-153 (*P* = 0.023) and Coll-524 (*P* = 0.004), and no notable difference from the tested transgenic strains. Further inoculation assays were conducted on *Capsicum annuum* cv. Fushimi-amanaga. Strains generated from transformation VII-X produced larger lesion sizes than wild-type Coll-365. The mean lesion sizes were increased between 49 and 400%. The lesion size increase level was similar for each transgenic strain in two independent experiments **(Table 9)**. Transgenic strains of transformation VI and X were further inoculated on *Capsicum annuum* cv. Groupzest and they caused significant lesion size increase as well (**S6 Table**).

**Figure 10.**
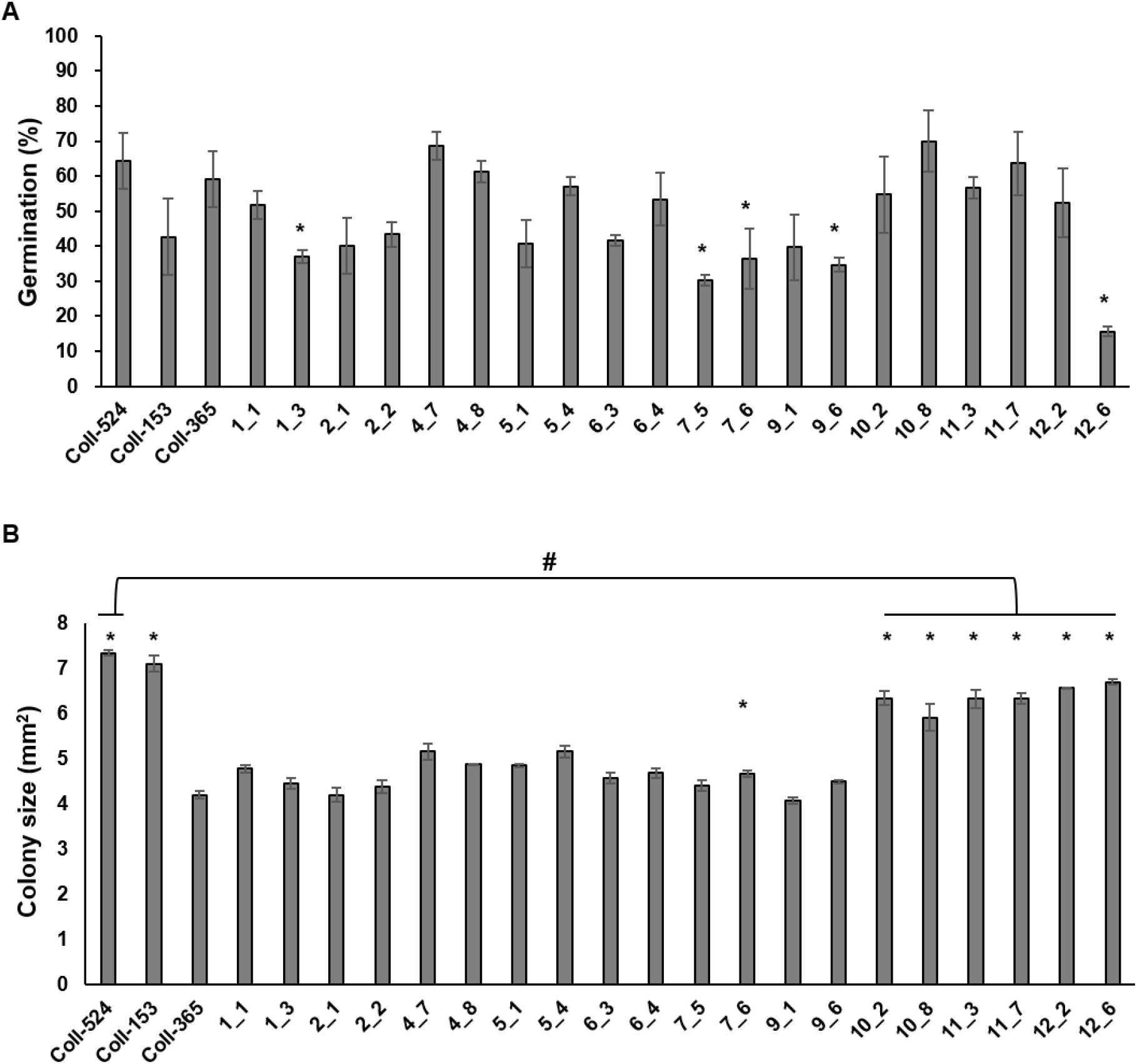
Spore germination (A) and mycelial growth of strains Coll-524, Coll-153 and Coll-365 and the transgenic strains of Coll-365 (B). The significant difference (*P* < 0.5) between Coll-365 and other strains was indicated with a star when *P* < 0.5 appeared in two independent experiments. The significant difference between Coll-524 and six transgenic strains of Coll-365 was indicated with a hash in panel B.

**Table 8.** Genetic transformation and phenotyping in strain Coll-365.

| Transformation | Gene(s)<sup>a</sup> | BLAST result | ORF length (bp) | Transgenic strains used for phenotyping | Functions |
| --- | --- | --- | --- | --- | --- |
| I | 11975 | hypothetical protein | 683 | 1-1, 1-3 | ND<sup>b</sup> |
| II | 14591 | probable laccase precursor | 2264 | 2-1, 2-2 | ND |
| III | 14976 | endo-β-1,3-glucanase | 2202 | 6-3, 6-4 | ND |
| IV | 15215 | LysM domain-containing protein | 1599 | 7-5, 7-6 | Inhibited germination |
| V | 15022 | bZIP transcription factor | 871 | 4-7, 4-8 | ND |
| VI | 15029 | C6 zinc finger domain-containing protein | 1050 | 5-1, 5-4 | ND |
| VII | 15022<br>15029 |  |  | 9-1, 9-6 | Enhanced virulence |
| VIII | 15019 | related to phospholipase a-2-activating protein | 2425 | 10-2, 10-8 | Enhanced growth and virulence |
| IX | 15022<br>15029<br>15019 |  |  | 11-1, 11-7 | Enhanced growth and virulence |
| X | 15022<br>15029<br>15019 |  |  | 12-2, 12-6 | Enhanced growth and virulence |
|  | 15003 | WD domain<br>containing protein | 1387 |  |  |

<sup>a</sup>, Gene(s) of Coll-524 used to be transferred into Coll-365 in each transformation experiment.
<sup>b</sup>, ND: non-detected; UD: undetermined.

**Table 9.** Pathogenicity assay of gene transformation strains (transgenic) and wild-type strain Coll-365 (WT) on fruits of *Capsicum annuum* cv. Fushimi-amanaga.

| Exp. | Transgenic strains | N<sup>a</sup> | Mean of lesion size |  | P | Lesion size increased |  |
| --- | --- | --- | --- | --- | --- | --- | --- |
|  |  |  | (mm<sup>2</sup>) |  |  | (%) |  |
|  |  |  | WT | Transgenic |  | Individual Exp. | Means of Exp. 1 & 2 |
| 1 | 4_7 | 5 | 16.50 | 19.43 | 0.2467 | 18 | 2 |
|  | 4_8 | 6 | 29.73 | 23.07 | 0.1429 | -22 | -27 |
|  | 9_1 | 4 | 14.18 | 26.00 | 0.0102 | 83 | 284 |
|  | 9_6 | 4 | 18.60 | 33.90 | 0.0916 | 82 | 71 |
|  | 10_2 | 7 | 12.90 | 60.48 | 0.0000 | 369 | 274 |
|  | 10_8 | 6 | 21.95 | 37.42 | 0.0814 | 70 | 49 |
|  | 11_3 | 8 | 13.26 | 49.43 | 0.0000 | 273 | 221 |
|  | 11_7 | 9 | 24.88 | 31.04 | 0.0364 | 25 | 54 |
|  | 12_2 | 9 | 10.11 | 58.87 | 0.0000 | 482 | 400 |
|  | 12_6 | 8 | 23.97 | 49.00 | 0.0000 | 104 | 123 |
| 2 | 4_7 | 7 | 22.72 | 19.67 | 0.5799 | -13 | 2 |
|  | 4_8 | 6 | 33.73 | 23.20 | 0.1201 | -31 | -27 |
|  | 9_1 | 5 | 5.63 | 32.88 | 0.0001 | 484 | 284 |
|  | 9_6 | 5 | 19.00 | 30.40 | 0.0665 | 60 | 71 |
|  | 10_2 | 7 | 23.60 | 65.87 | 0.0001 | 179 | 274 |
|  | 10_8 | 7 | 24.80 | 31.53 | 0.2146 | 27 | 49 |
|  | 11_3 | 8 | 13.04 | 35.13 | 0.0000 | 169 | 221 |
|  | 11_7 | 6 | 14.64 | 26.86 | 0.0269 | 83 | 54 |
|  | 12_2 | 8 | 11.00 | 45.89 | 0.0000 | 317 | 400 |
|  | 12_6 | 7 | 12.91 | 31.26 | 0.0015 | 142 | 123 |

<sup>a</sup>, N indicates the numbers of fruits used in each inoculation.

## Discussion

*Colletotrichum* species can cause great economic loss to various crops. Among the *Colletotrichum*, more than 30 species have been documented to cause chili anthracnose disease, constituting a major limitation to chili pepper production in tropical and subtropical regions [36]. In Taiwan and other Asia countries, *C. scovillei* attacks chili fruits [4], but its interactions with hosts at the molecular genetic level remain to be examined. In this study, we focused on genomic comparisons of the three *C. scovillei* strains and combined these with genetic approaches to identify genes involved in fungal growth and virulence. We have provided the genome sequences of the three strains with gene functional annotation for *C. scovillei*. We have setup a simple mathematical method to search for ORF variations between Coll-153 or Coll-365 and Coll-524 and successfully identified DNA fragments containing genes involved in the defects in growth and virulence of Coll-365. Moreover, by genetic assay we have demonstrated four genes that have functions in germination, growth and/or virulence of *C. scovillei*.

Our data suggested that the three strains all belong to the acutatum complex and are members of *C. scovillei* because they were grouped in the same clade of *C. scovillei* CBS 126529, the holotype strain of *C. scovillei* [2]. The data were consistent with a previous study that showed that Coll-524 and Coll-153 are in the *C. scovillei* clade of acutatum species complex [4].

For closely related species, de-novo assembly by reference-guided or mapping through a well-sequenced species is efficient, and can often improve the completeness of the genome sequence [37]. Our data revealed the major variation of Coll-153 and Coll-365 was the result of sequence removal. Genetic variations can be caused by three different events, local nucleotide sequence changes, intragenomic rearrangement of DNA segments and the acquisition of a foreign DNA segment by horizontal gene transfer [38]. The transposable element is one of the major factors leading to variations in fungi. In *C. higginsianum*, two closely related strains carry large-scale rearrangements and strain-specific regions that are frequently associated with transposable elements [39]. In the two *C. higginsianum* strains, the gene-sparse regions are transposable element-dense regions that have more effector candidate genes, while gene-dense regions are transposable element-sparse regions harboring conserved genes [39]. *C. higginsianum* and other eukaryotic plant pathogens, such as *Phytophthora infestans* and *Leptosphaeria maculans* have been referred to as “two-speed genomes” as these genomes have a compartmentalized genome structure to protect housekeeping genes from the deleterious effects of transposable elements and to provide rapid evolution of effector genes [40, 41]. In the genomes of Coll-524, 153 and 365, we did not find a relationship between the density of transposable elements and effector genes or housekeeping genes. Coll-524 and *C. higginsianum* IMI 349063 have similar genome sizes but *C. higginsianum* IMI 349063 carries nearly three times the number of transposable element genes to Coll-524 (S7 Table). Moreover, compared with *C. higginsianum* IMI 349063, Coll-524 was found to be unlikely to have sequence structures like mini chromosomes which harbor high density transposable elements with over 40% sequences encoding transposable elements [42].

We designed a simple mathematical method to identify ORF-variations. Applying this method, a 34-kb fragment (Fig 5) containing 14 genes that exist in the Coll-524 genome but are almost completely lost in Coll-365 were identified. The loss of the 14 genes in Coll-365 is likely caused by DNA deletion that might have resulted from DNA rearrangements. DNA rearrangement frequently occurs in fungi via the parasexual cycle that leads to recombination and chromosome gain or loss [43]. It is possible that Coll-524 gained part or all of these genes by DNA rearrangement during the parasexual cycle in Coll-153 or Coll-365. Coll-365 and Coll-153 were collected 4 years earlier than Coll-524 from the fields by the ACRDC-the World Vegetable Center [9]. Coll-153 and Coll-365 are grouped in the CA1 pathotype and Coll-524 is a member of the CA2 pathotype. CA2 has higher virulence than CA1 and has replaced CA1 to become dominant in Taiwan [11, 12]. Thus, Coll-524 might have arisen evolutionarily from the CA1 pathotype members such as Coll-153 and Coll-365 by gaining genes horizontally or via gene arrangement. They might be evolutionarily developed from different branches of *C. scovillei*.

Gene family expansions and contractions are related to the changing of host range and virulence of plant pathogens [17, 44]. A set of 33 effectors was completely lost in Coll-365, but existed in two closely related chili pepper pathogens *C. acutatum* strain 1 and *C. scovillei* strain TJNH1. This suggests that the 33 effectors might play a role in the virulence of Coll-524 to chili pepper. However, when one of the effectors was transferred to Coll-365, it did not promote the virulence of the transgenic strains on chili pepper fruit, suggesting that multiple effectors may work together to affect fungal virulence [45].

CAZyme genes are potentially involved in fungal growth and colonization of host tissues. Coll-524, Coll-153 and Coll-365 have 661-663 CAZyme genes. In a study that compared the protein family encoded by *Colletotrichum* species and others, CAZyme GH43 and S10 protease were found to be duplicates with specific genes within the respective lineages in the wide host range pathogens acutatum and gloeosporioides species complex [17]. Research on Magnaporthaceae specific clusters showed CAZyme gene families may contribute to speciation [46]. Transgenic strains carrying endo-beta-1,3-glucanase or laccase did not result in a gain of function of growth and virulence of chili pepper. Endo-beta-1,3-glucanase is expected to play a key role in cell wall softening in ascomycetes. The endo β-1,3-glucanase in *Schizosaccharomyces pombe, Candia albicans* and *Saccharomyces cerevisiae* has been showed to be required for dissolution of the primary septum to allow cell separation [47–49]. Fungal laccase can detoxify phenolic pollutants and has a role in detoxifying host defense phenolic compounds, such as capsaicin in chili pepper [50]. Coll-524 has 5 copies of laccase-related genes and two endo-beta-1,3-glucanase and one endo-beta-1,3(4)-glucanase. A single laccase or endo-beta-1,3-glucase gene may not be able to contribute the function of growth or virulence of Coll-365.

Eight genes were selected for gene transformation in Coll-365. Three genes 11975, 14591 and 14976 did not show a role in germination, growth and virulence of Coll-365 after transforming into Coll-365. The effector gene 11975, which encodes a hypothetical protein, was expressed strongly and specifically at the early infection stage, while laccase gene 14591 was expressed highly at all infection stages but not in axenic cultures. Endo-beta-1,3-glucanase gene 14976 was highly expressed in axenic cultures and at the late infection stage. Gene 15215 encoding a LysM domain-containing protein was expressed at extremely high levels in axenic culture and at all infection stages. The transformation of gene 15215 to Coll-365 showed no significant influence on fungal growth and virulence but inhibited spore germination ability. LysM domain-containing protein can bind to the fungal cell wall and protect from cell wall-degrading enzyme digestion and functions in ecological niche competition and prevention from recognition by the host to trigger host immunity [51]. The LysM domain-containing protein TAL6 of *Trichoderma atroviride* was shown to specifically inhibit spore germination of *Trichoderma* species and was suggested to be involved in the self-signaling process but not fungal-plant interactions [52]. *Colletotrichum* species can self-inhibit spore germination by producing self-inhibitors [53]. Whether this LysM domain-containing protein encoded by gene 15215 is involved in germination self-inhibition under high spore concentration and pathogenicity in Coll-524 needs to be verified in the future.

Four genes (15022, 15029, 15019, 15003) located within the 34-kb fragment at scaffold 17 were transferred into Coll-365 as individual genes or gene sets since they are closely linked. Gene 15022 had a high expression level in axenic cultures and all infection stages. The other three genes were expressed in axenic cultures and the late infection stage, but genes 15019 and 15029 also had relatively high expression in the cuticle infection stage. These four genes did not influence the germination and appressorium formation of Coll-365. However, transgenic strains carrying one gene 15019, three genes 15022, 15029, and 15019, or four genes 15022, 15029, 15019, and 15003 have similar colony sizes but are larger than the wild-type Coll-365, indicating that gene 15019 contributes to the growth enhancement of Coll-365. Gene 15022 is a bZIP transcription factor and might be highly related to pathogenicity [54]. The opposite direction gene 15029, 1,313 bp away from gene 15022 encodes a C6 zinc finger domain-containing protein. The C6 zinc finger proteins are strictly fungal proteins and involved in fungal-host interactions [55]. Gene 15022 or 15029 alone did not affect the virulence of Coll-524. However, transgenic strains carrying the two genes can enhance the virulence of Coll-365, indicating that the co-existence of the two genes together might contribute to the virulence enhancement of Coll-365.

Gene 15019 encodes a protein related to phospholipase A2-activation, while gene 15003 encodes a WD domain-containing protein. Transgenic strains carrying one gene 15019, three genes 15022, 15029, and 15019, or four genes 15022, 15029, 15019, and 15003 enhance the virulence of Coll-365, indicating gene 15019 might be the major contributor to the virulence enhancement of Coll-365. Moreover, transgenic strains carrying genes 15022, 15029, and 15019, or genes 15022, 15029, 15019, and 15003 have a smaller *P* value than transgenic strains carrying gene 15019 only (Table 9), which is consistent with the virulence contribution by gene 15022 and 15029 as mentioned above. Gene 15019 encodes a protein related to phospholipase A2-activation. Phospholipase A2-activating protein has been relatively intensively studied in humans and is involved in apoptosis and tumor regression; however, there is only a single study related to its functional characterization in fungi [56, 57]. The *Magnaporthe royzae Moplaa* gene encodes a phospholipase A2-activating protein and the deletion of this gene results in the reduction of fungal growth and pathogenicity [56].

In conclusion, in this study, four genes were identified which function in germination, growth and/or virulence of the chili pepper pathogen *C. scovillei.* Gene 15215, encoding a LysM domain-containing protein, reduce the germination of Coll-365. Genes 15029 and 15022, encoding a C6 zinc finger domain-containing protein and a bZIP transcription factor, together enhance the virulence of Coll-365. Gene 15019, encoding a protein related to phospholipase A2-activation, enhances the growth and virulence of Coll-365. In addition, the 34-kb fragment in scaffold 17 contributes a lot to the defects in growth and virulence in Coll-365 because three genes 15019, 15022 and 15029 are located on this fragment. Interestingly, we also identified 33 effectors lacking in Coll-153 and Coll-365 which may be involved in the full virulence of Coll-524 on chili pepper. Future study will focus on the 33 effectors and other effectors that may have led to Coll-524 becoming a dominant and virulent pathogen.

## Materials and Methods

### Fungal strains and culture conditions

Three *Colletotrichum* strains Coll-524, Coll-153 and Coll-365 were obtained from the AVRDC-The World Vegetable Center (Tainan, Taiwan) as mentioned in a previous study [9]. For sporulation, fungal strains were cultured on MS agar medium (0.1% yeast extract, 0.1% peptone, 1% sucrose, 0.25% MgSO_4_•7H_2_O, 0.27% KH_2_PO_4_ and 1.5% agar) for 6 days at 25°C with a 12 h light/dark cycle. Spores were collected and used for general cultivation, assays of germination and appressorium formation, morphological examination, growth, and pathogenicity assay.

### Genomic DNA extraction and library preparation for sequencing

Spores were inoculated into a 500 mL flask containing 100 mL MS liquid medium (1 x 10^5^ spores/ml) at 25°C with shaking at 200 rpm for 1 day. The young hyphae were collected by filtrating with a layer of miracloth. The DNA was extracted from the young hyphae using the CTAB extraction method [58]. DNA purity and concentration were determined by Nanodrop 2000 (Thermo Fisher) and Qubit (Invitrogen) measurements. Libraries were generated from 2-5 μg of genomic DNA using the TruSeq 2 library preparation kit (Illumina, USA). Two types of libraries were prepared, short-fragment library (0.3 kb) for paired-end sequencing and long-fragment library (3 kb and 8 kb) for mate-pair sequencing. DNA fragmentation was made by shearing using an M220 Focused-Ultrasonicator (Covaris, USA) and the fragments with 296-300 bp, 3 kb or 8 kb were selected by gel-cutting. Without the 120 bp adaptor, the remaining 176-180 bp DNA fragments may have 20% overlap with 100 bp read length for paired-end sequencing. Long-fragment libraries, 3 kb and 8 kb, were prepared for Coll-524 only.

### Genome sequencing and assembly

The Illumina HiSeq 2500 platform was used to sequence the genome of Coll-524 with 100 bp read length for paired-end and 3 kb mate pair libraries, and 125 bp read length for 8 kb mate pair library. The genomes of Coll-153 and Coll-365 were sequenced by Illumina Genome Analyzer with 150 bp read length. The raw reads were trimmed by Trimmomatic 0.39 to remove linkers and adaptors. Those refined reads of Coll-524 were further assembled by the ALLPATHS-LG *de novo* assembly program [30]. The genome sequences of Coll-524 have been deposited in NCBI with accession numbers JAESDM010000001-JAESDM010000054.

Genome assembly of Coll-153 and Coll-365 was achieved by using the “Map Reads to contigs” function in CLC Genomic Workbench v.9.5.1. All trimmed reads were used to map to the Coll-524 genome under default settings. The consensus mapped sequences were extracted and joined to form a scaffold by the “Extract Consensus Sequence” function. Unmapped reads were collected and assembled by using the “De Novo Assembly” function in CLC software. The assembled contigs with sizes larger than 5 kb were collected and counted as the genome sequences of Coll-153 or Coll-365. The genome sequences of Coll-153 and Coll-365 have been deposited in NCBI with accession numbers JAESDN010000001-JAESDN010000059 for Coll-153 and JAESDO010000001-JAESDO010000059 for Coll-365.The genome completeness of the three strains were assessed using Benchmarking Universal Single-Copy Orthologs (BUSCO v.3.0) and the sordariomyceta_odb10 dataset [32, 59].

### Multi-locus phylogenetic analysis

The DNA sequences of five marker genes (ACT1, CHS1, GAPDH, ITS and TUB2) were used for phylogenetic analyses [1]. The sequences of the five genes from the three strains, 170 other *Colletotrichum* strains and an outgroup species, *Monilochaetes infuscans*, were obtained and used in the assay. Among them, the five marker genes of 108 *Colletotrichum* species and *M. infuscans* were obtained from Q-bank database (https://qbank.eppo.int/fungi/) according to the GenBank accession numbers listed in a publication of Cannon *et al.* [1], while sequences from 62 *Colletotrichum* strains were obtained by downloading whole genome sequences from NCBI FTP and then identifying the five gene sequences by using CLC Genomic Workbench v.9.5.1. The information for all the genes sequences is provided in **S1 Table**. All collected sequences were aligned by MAFFT v.7 online version [60] with default settings and then trimmed by trimAl 1.4.1 [61] with automated1 setting. Five marker genes were concatenated in each strain. Bayesian inference of all the concatenated sequences was analyzed by MrBayes 3.2.7 [62]. The Markov Chain Monte Carlo (MCMC) chains were set for 5,000,000 generations. Sample frequency was set for every 100 generations. Phylogenetic tree of the 174 strains were generated by FigTree v.1.4.3 [63].

### Genome annotation

Gene annotation was conducted using MAKER (v.2.31.10) [31] pipeline with default settings (Augustus, PJ7, RNAseq reads). The genome of Coll-524 was used for gene annotation. The assembled contigs from Coll-524 RNAseq data was provided for EST evidence. The RNAseq data of Coll-524 included 10 sets of RNAseq data, which were infected purified cuticle layer, axenic 18 h hyphal culture, and 8 sets of infected chili pepper fruits at different infection time points (unpublished). The CDS and protein sequence of *Colletotrichum fioriniae* PJ7 (PRJNA244481) was provided to MAKER for the “EST evidence” and “Protein Homology Evidence” functions respectively. The gene annotation of Coll-153 and Coll-365 was conducted using the same method as that for Coll-524. The CDS region of each gene was extracted from the genome sequence according to the GFF file which was originated from MAKER by GffRead [64].

### Gene functional analysis, clustering and orthology analysis

The genes of Coll-524, Coll-153 and Coll-365 were searched with nr database by using BLASTx (BLAST 2.2.29+) to obtain the information about gene function. To find effector candidates, protein sequences of the three strains were analyzed by EffectorP 2.0 [65]. Carbohydrate active enzymes (CAZymes) candidates were searched by dbCAN2 meta server [66]. Candidate gene selection followed the recommendation setting provided by dbCAN2 meta server. Genes related to secondary metabolism were predicted using the Secondary Metabolite Unique Regions Finder (SMURF) database [67]. The Pathogen-Host Interactions database (PHI-base) v.4.10 was downloaded from PHI-base website. All genes of the three strains were used to blast against PHI-base using CLC Genomic Workbench [68]. Genes with E-values lower than 10 were selected according to the default setting of the PHIB-BLAST web server. The protein sequences of all annotated genes of Coll-153, Coll-365 and Coll-524 were used for ortholog analysis with the default settings of OrthoFinder [33]. After the Orthofinder analysis, single-copy orthogroups were generated directly. Other analyses of orthologues, including multi-copy orthogroups with the same gene numbers, multi-copy orthogroups with different gene numbers, orthogroups in every two strains, strain-specific orthologues, and unassigned genes, were further classified by EXCEL. These unassigned genes were considered as “strain specific genes” together with genes in strain specific orthogroup. Genes of Coll-524 with extra copy number in comparison with the other two strains in the orthogroup, and genes analyzed to be Coll-524 strain specific genes were further considered as “Coll-524 extra genes”.

### Identification of microsatellites, repeats and transposon elements

MISA 2.0 was used for microsatellite analysis [34] and transposon elements were analyzed by Transposon PSI 2.2.26 (http://transposonpsi.sourceforge.net/). The transposon element sequences were acquired according to the GFF files provided by Transposon PSI. These sequences were further extracted to create a Transposon element database (TED). RepeatModeler was used for de novo genome-wide repeat family analysis [35]. The sequences of TcMar-Fot1 were further blasted with BLASTn to the NCBI database and the functional domains were analyzed using InterPro [69] to determine if DDE was present [70].

### Identification of genome sequence variations and ORF variations of Coll-153 and Coll-365

During the mapping of the Coll-153 or Coll-365 genome to the Coll-524 genome, data containing the variations between Coll-524 and the other 2 strains was generated with the “Extract Consensus Sequence” function. The data included four datasets, Misc. difference (MD), Insertion, Deletion and Removed, which are specified by CLC with default setting. Briefly, “Deletion” is for positions where a gap is called while the reference has a non-gap; “Insertion” is for positions where a non-gap is called while the reference has a gap; “Removal” is for positions where no reads are mapped to the reference; “Misc. difference” is for every position where the consensus is different from the reference. The MD dataset consisted of single nucleotide polymorphism (SNP) and polymorphism with more than one nucleotide, which was named as multiple nucleotide polymorphism (MNP) in this study. The four datasets were combined to generate the genome sequence variations and used for genome comparisons to identify ORF variations between Coll-524 and the other two strains as described in **Fig 4**. To simplify the analysis of ORF variations, the Removed datasets of Coll-153 and Coll-365 were combined. Briefly, all the sequences which existed in Coll-524 but removed from Coll-153 or Coll-365 were extracted from Coll-524 genome to create a “removed sequence database” (RSD).

ORF variation analysis was based on the calculations of physical position overlapping between each ORF and the four datasets of variations (MD-MNP, Insertion, Deletion and Removed). The GFF file generated by MAKER could provide the physical positions of mRNA, exon, 5′UTR, CDS, 3′UTR for each gene. A gene usually contains multiple CDSs in eukaryotes and all CDSs together within the gene is the complete coding sequence to translate a protein encoded by the gene. To analyze ORF variation of Coll-153 and Coll-365, the ORF position of each Coll-524 gene was generated by setting the first nucleotide of the first CDS as the beginning position and the last nucleotide of the last CDS as the end position.

To analyze the variation of each ORF between Coll-524 and the other two strains, firstly, all scaffolds of Coll-524 were added together one by one to form a single sequence. The physical positions of each ORF were modified to fit into the single sequence. The four datasets of variations (MD, Insertion, Deletion and Removed) generated by CLC were tagged with physical positions in each scaffold of Coll-524. Therefore, the second step was to edit the physical positions of each variation according to the single genome sequence of Coll-524. The position differences of each variation to the ORF were compared and calculated as indicated in **Fig 4C-G**. The physical position overlapping situation of each ORF and each variation was identified by the calculation formula listed in **Fig 4G**. An ORF overlapped with a variation indicated that this ORF varied between Coll-524 and the other strain carrying this variation. ORFs overlapped with variation were further called “ORF-V”.

### Screening of genes potentially involved in the variations of the three strains for functional verification in Coll-365

To select genes for genetic analysis to verify their functions on the defects of growth and virulence of Coll-365, four selection criteria were set for gene selection. First were ORFs predicted to be ORF-V. Second were “Coll-524 extra genes” according to the result of OrthoFinder analysis. Third were genes analyzed to belonging to effector, CAZyme, TF, SMURF, enzyme or PHI (E-value < 10e-50). Finally, were ORFs with <50% coverage to Coll-153 and/or 365 according to the blastn results.

Genes with the four criteria were analyzed by Venn Diagram (http://bioinformatics.psb.ugent.be/webtools/Venn/). Pathogen-host interaction related genes were selected with an E-value less than 10e-50. Genes with an E-value less than 10e-50 but greater than 10e-100 were considered to have almost identical sequences to the database according to the manual for CLC Genomics Workbench 9.5.

### Plasmid construction

Plasmids BsHR and BsBR were constructed and used in this research to clone the selected target gene(s) for transformation into Coll-365. pBsHR was constructed by cloning hygromycin resistance cassette (HygR) of pBHt2 [71] into pBluescript SK(+) [72] with PCR amplification, restriction enzyme digestion (EcoRV and SwaI, designed in the primers), and DNA ligation. Plasmid BsBR was modified from pBsHR by replacing (HygR) with bleomycin resistance gene (BleoR) of pAN8-1 [73]. The BleoR cassette was PCR amplified with specific primers containing EcoRV and SwaI restriction sites and then cloned into EcoRV- and SwaI-digested pBsHR. To clone the target genes from Coll-524, a target gene containing an approximate 1-kb promoter and 0.5-kb terminator was amplified with a high-fidelity DNA polymerase KOD-plus-Neo (TOYOBO, Osaka, Japan) and ligated to EcoRV or SwaI digested and Shrimp Alkaline Phosphatase (rSAP) treated pBsHR. If double gene-transformation was needed, a second target gene was cloned in to pBsBR for protoplast transformation and phleomycin was used as the selection antibiotic [74]. For gene-transformation VII, the two genes were closely linked and were amplified together with one primer set by PCR and transferred to Coll-365 by single transformation. Gene-transformation IX was completed with two transformations by transferring gene 15019 into a transgenic strain of gene-transformation VII. The gene-transformation X was conducted by transferring a PCR fragment containing 15019 and 15003 into a transgenic strain of gene-transformation VII. The gene locations of 15019, 15003, 15022 and 15029 are indicated in **Fig 5**.

### Protoplast transformation

Polyethylene glycol (PEG)/Ca^+2^-mediated protoplast transformation was used to transfer the target gene to Coll-365. Spores collected from a 6-day MS agar medium were inoculated into a 500-ml flask containing 400 ml MS liquid medium and a stirrer bar, and cultured for 16 h under 25°C at low speed to prevent the attachment of spores to the flask surface. The young hyphae were collected by filtering through a layer of miracloth and washed with 100 ml sterilized water and 100 ml wash buffer (1 M NaCl and 10 mM CaCl_2_) under suction. The washed hyphae (250 mg) were then resuspended in a 50-ml flask containing 10 ml osmotic buffer (10 mM Na_2_HPO_4_, pH 5.8, 20 mM CaCl_2_, and 1.2 M NaCl) with 90 mg lysing enzymes (Sigma, L1412), and incubated in an orbital shaker with 150 rpm at 30°C for 6 h. The undigested hyphae were removed by filtering through miracloth and the protoplasts were collected by centrifugation at 1500 g, 4°C for 10 min. The protoplasts were resuspended by adding 100 μl mixture of four parts of STC buffer (1.2 M sorbitol, 10 mM Tris-HCl, pH 7.5, and 10 mM CaCl_2_) and one part of PEG [50% (w/v) polyethylene glycol 3350, 10 mM Tris-HCl, pH 7.5, and 10 mM CaCl_2_]. For transformation, 20 μl of 20 μg plasmid DNA was added to a tube containing 100 μl protoplast suspension with gentle mixing and the mixture was placed in ice for 20 min. PEG was added into the DNA-protoplast solution four times with different volumes. The adding steps were performed by following a previous description [75] with slight modifications. Briefly, 20, 80, 300 and 600 μl PEG were added into protoplast suspension step by step. After each addition, the mixture was gently mixed and left to stand for 3 min, and then left to stand a further 20 min after the last addition at room temperature. After the 20 min incubation, 3 ml regenerate buffer [4 mM Ca(NO_3_)_2_·4H_2_0, 1.5 mM KH_2_PO_4_, 1 mM MgSO_4_·7H_2_O, 2.5 mM NaCl, 60 mM glucose, and 1 M sucrose] was added into the PEG-DNA-protoplast mixture and cultured at 25°C for 16 h with 100 rpm shaking. The protoplasts were collected with centrifugation at 25°C, 1800 g for 10 min. A total of 300 μl regenerated protoplasts were obtained and then evenly spread on three regeneration agar medium plates. After incubation for 6-8 days at 25°C, the transformants were isolated, single spore purification and PCR verification for the target gene insertion.

### PCR analysis for gene loss in Coll-153 and Coll-365 and verification of gene transformation in Coll-365 transformants

Eight genes were selected for functional assay in Coll-365 as listed in **Table 8**. PCR assays were used to confirm the loss of the eight genes in Coll-365 using specific primer sets as listed in **S8 Table**. Genomic DNA was extracted as described above and tubulin gene (Accession number:MW073123) was used as the control for PCR assay. To verify gene transformation of transgenic Coll-365 strains, regular PCR and RT-PCR assays were performed. Regular PCR using genomic DNA as template to amplify the transgenic gene was used for all transgenic strains. For transgenic strains carrying more than one transgene, RT-PCR assays were conducted in addition to the regular PCR. Primers used in theses assays are listed in **Table S6**. For RNA extraction, spores were inoculated into a 50 mL flask containing 20 mL MS liquid medium (1 x 10^5^ spores/ml) at 25°C with shaking at 150 rpm for 2 days. The hyphae were collected by filtrating with a layer of miracloth. The RNA was extracted with Trizol reagent (Invitrogen). RNA purity and concentration were determined and cDNA was synthesized using 5 μg RNA and the MMLV reverse transcription kit (Invitrogen). PCR was performed using 20 μl reaction volume contained 1 × PCR Buffer, 0.2 mM dNTP, 0.4 μM primers, 1 U Blend Taq plus (TOYOBO), and 1 μl of template DNA. PCR reaction was performed as follows: 94°C for 2 min and 25 cycles of 94°C for 30 sec, 60°C for 30 sec, and 72°C for 3 min. The PCR products were analyzed by agarose gel electrophoresis.

### Fungal growth, spore germination and appressorium formation assays

Fungal growth was assayed by inoculating spore suspension on the center of three different agar media. Spore germination and appressorium formation were detected in 96-well plates. Briefly, spores were collected from MS agar medium and the concentration was adjusted to 2 × 10^5^ spores/ml by sterilized water. For growth assay, a 5-μl, spore suspension containing 500 spores was dropped on the center of MS agar medium plate and incubated for 6 day at 25°C with a 12 h light/dark light cycle. Colony sizes were measured by using ImageJ 1.53a [76]. To detect germination and appressorium formation, 80 μl of spore suspension was added into each well of a 96-well plate and incubated at 25°C with 12 h light/dark light cycle for 8 h. To count the numbers of germinated spores and appressorium, the 96-well plate was placed upside down and examined using a light microscope. All the experiments mentioned above were conducted at least two times with three replicates for each strain at each time. The statistical significance of the data was determined by One-way ANOVA at *P* < 0.05 using Statistical Package for the Social Sciences software, version 20 (IBM SPSS software).

### Pathogenicity assay

Pepper fruits were used for pathogenicity assay, including *Capsicum annuum* cv. Hero, Fushimi-amanaga and Groupzest. The fresh harvested fruits were washed with water and surface sterilized with 0.5% bleach and left to dry overnight. Fungal spores were collected from MS agar medium and the concentration was adjusted to 2 × 10^5^ spores/ml. Spore suspension was dropped (5 μl/drop) on the fruit surface with dual inoculation, in which the wild type strain was inoculated on one side of the fruit and a transgenic strain was inoculated on the other site of the same fruit. The fruits were incubated in a growth chamber at 25°C with 12 h light/dark light cycle. At least three fruits having 3-5 inoculation sites were used for each strain in each inoculation. Lesion sizes were measured with ImageJ and the statistical analysis was performed by paired *t*-test at *P* < 0.05 using Statistical Package for the Social Sciences software, version 20 (IBM SPSS software).

## AUTHOR CONTRIBUTIONS

DKH, SCC, MHL, and MCS contributed to the design of the experiments. SCC designed and prepared the DNA and RNA materials for sequencing. DKH performed all bioinformatic analyses and gene transformation as well as functional characterizations. YTC and CYC assisted in sequence assembly and annotation. MYL performed NGS sequencing. MHL and MCS supervised the experiments. DKH, MHL, and MCS wrote the manuscript.

## ACKNOWLEDGMENTS

The authors would like to thank Academia Sinica High throughput Genomics Core for NGS sequencing services, and Ms. Miranda Loney for English editing.

## SUPPLEMENTARY MATERIAL

The Supplementary Material for this article can be found online at:

## Supporting information

### Supporting Tables

**S1Table. Information on the five maker genes (ACT1, CHS1, GAPDH, ITS and TUB2) used in the phylogenetic analysis in Fig 1 and S1 and S2 Figs.**

**S2 Table. Functional category statistics of the three *Colletotrichum scovillei* genomes.**

**S3 Table. Functional category statistics of the multi copy orthogroups with different gene numbers for the three *Colletotrichum scovillei* genomes.**

**S4 Table. Repeat sequence analysis with MISA for microsatellite compositions in the three strains.**

**S5 Table. The distribution of the 59 genes clustered at different scaffolds in Coll-524.** The numbers of genes absent in both strains Coll-153 and Coll-365, and in strain Coll-365 only, are presented.

**S6 Table. Pathogenicity assay of transgenic strains and wild-type strain Coll-365 (WT) on fruits of *Capsicum annuum* cv. Groupzest by pair inoculation.**

**S7 Table. TransposonPSI analysis results of three *Colletotrichum scovillei* strains and *C. higginsianum* IMI 349063.**

**S8 Table. Primers used in this study.**

### Supporting Figures

**S1 Fig. Phylogenetic tree of 173 *Colletotrichum* strains and one *Monilochaetes infuscans* strain showing the phylogenetic relationship of the three *Colletotrichum* strains (Coll-524, Coll-153 and Coll-365) and other *Colletotrichum* species.** The phylogeny was constructed by using DNA sequences of five markers (ACT1, CHS1, GAPDH, ITS and TUB2) according to Cannon et al. (2012) with additional 62 *Colletotrichum* strains from NCBI as indicated in Table S1. The 62 strains from NCBI was marked with “*”. Values at the nodes are Bayesian posterior probability values. Strains belong to various *Colletotrichum* species complex were indicated with different color boxes.

**S2 Fig. The original host and phylogenetic relationship of strains in *Colletotrichum acutatum* species complex. This tree was part of the phylogenetic tree in S1 Fig.** The original host of each strain was indicated in the right side behind each strain. The clade containing Coll-524, 153 and 365 was highlighted with yellow color.

**S3 Fig. Drug metabolism – cytochrome P460 KEGG pathway (A) and the gene number variations of monooxygenase among the three strains (B).** (A) The drug metabolism KEGG map shown here is a partial map of map00982. (B) The gene number differences of monooxygenase among the three strains were based on the investigation of scaffolds 17, 19, 20 and 22. The positions of monooxygenase in the drug metabolism – cytochrome P450 pathway are indicated with red circles in panel A.

**S4 Fig. Purine metabolism KEGG pathway (A) and the gene number variations of adenylpyrophosphatase among the three strains (B).** The position of the adenylpyrophosphatase in the purine metabolism pathway is indicated with a red circle in panel A.

**S5 Fig. The thiamine biosynthesis KEGG pathway (A) and the gene number variations of phosphatase among the three strains (B).** The position of the phosphatasease in the thiamine biosynthesis pathway is indicated with a red circle in panel A.

**S6 Fig. Phenylpropanoid biosynthesis KEGG pathway (A) and the gene number variations of lactoperoxidase among the three strains (B).** The positions of the lactoperoxidase in phenylpropanoid biosynthesis pathway are indicated with red circles in panel A.

**S7 Fig. N-glycan biosynthesis KEGG pathways (A) and the gene number variations of alpha-1,6-mannosyltransferase among the three strains (B).** (A) The N-glycan biosynthesis KEGG map shown here is a partial map of map00513. (B) The gene number differences of 1,6-mannosyltransferase among the three strains were based on the investigation of scaffolds 17, 19, 20 and 22. The positions of the 1,6-mannosyltransferase in the purine metabolism pathway are indicated with red boxes and lines in panel A.

**S8 Fig. The complete Coll-Fot1 (1891 bp) of strain Coll-524 consists of a coding region and two inverted terminal repeats (ITRs).** The DDE_1 containing region provided by TransposonPSI is highlighted with a green arrow. The coding region of 545 aa sequence was identified by SnapGene and is highlighted with a light blue arrow. The yellow bar indicates the DDE_1 domain resulted from InterPro analysis. Two ITRs with 55 bp in length are provided by RepeatModeler and is highlighted as a blue arrow.

**S9 Fig. Verification of gene presence in strains Coll-524, 153 and 365 by PCR (A), and confirmation of transgenic strains with regular PCR (B) and RT-PCR (C) for target gene transformation in strain Coll-365.**

